# Beat-locked ATP microdomains in the sinoatrial node map a Ca^2+^-timed energetic hierarchy and regional pacemaker roles

**DOI:** 10.1101/2025.08.18.670947

**Authors:** Manuel F. Muñoz, Collin Matsumoto, Paula Rhana, Declan Manning, Geoanna Bautista, Daniel M. Collier, L. Fernando Santana

## Abstract

Pacemaker myocytes of the sinoatrial (SA) node initiate each heartbeat through coupled voltage and Ca^2+^ oscillators, but whether ATP supply is regulated beat-by-beat in these cells remains unclear. Using genetically encoded sensors targeted to the cytosol and mitochondria, we tracked beat-resolved ATP dynamics in intact mouse SA node and isolated myocytes. Cytosolic ATP rose transiently with each Ca^2+^ transient and segregated into high- and low-gain phenotypes defined by the Ca^2+^–ATP coupling slope. Mitochondrial ATP flux adopted two stereotyped waveforms—Mode 1 “gains” and Mode 2 “dips.” Within Mode 1 cells, ATP gains mirrored the cytosolic high/low-gain dichotomy; Mode 2 dips scaled linearly with Ca^2+^ load and predominated in slower-firing cells. High-gain/Mode 1 phenotypes localized to superior regions and low-gain/Mode 2 to inferior regions, paralleling gradients in rate, mitochondrial volume, and capillary density. Mechanistic dissection placed sarcoplasmic reticulum (SR) Ca^2+^ release upstream of ATP production, showing that Ca^2+^ triggers metabolic transients while membrane voltage primarily modulates their frequency. Inhibiting mitochondrial Ca^2+^ uptake and adenine nucleotide exchange eliminated beat-locked mito- and cyto-ATP signals, indicating that the mitochondrial Ca^2+^ uniporter (MCU)–adenine nucleotide translocase (ANT) machinery couples Ca^2+^ release to ATP fluctuations. Mode 2 recovery kinetics indicate slower ATP replenishment, which would favor low-frequency, fluctuation-rich firing in a subset of cells. Together, these findings reveal beat-locked metabolic microdomains in which Ca^2+^ transients time oxidative phosphorylation under a local O_2_ ceiling, unifying vascular architecture, mitochondrial organization, and Ca^2+^ signaling to match energy supply to excitability. This energetic hierarchy helps explain why some pacemaking myocytes are more likely to set the rate, whereas others may widen the bandwidth.

**Summary:** Beat-locked cytosolic and mitochondrial ATP transients in SA-node myocytes sort into high-gain, low-gain, or consumption-dominant modes aligned with superior–inferior vascular–mitochondrial gradients. This energetic hierarchy lets high-gain cells set fast rates while low-gain/dip cells stabilize slow rhythms, broadening operating range but capping maximal bandwidth.

## Introduction

The heartbeat originates in the sinoatrial (SA) node, a crescent-shaped pacemaker region embedded at the junction of the superior vena cava and right atrium. Within the node, clusters of pacemaker myocytes fire spontaneous action potentials that propagate through gap junctions to neighboring SA node myocytes and, ultimately, the surrounding myocardium.

Pacemaking emerges from the concerted operation of a “membrane clock,” composed of voltage-gated ion channels, including hyperpolarization-activated, cyclic nucleotide-gated channels (HCN2/HCN4) and Ca^2+^ channels (Ca_V_3.1, Ca_V_1.3, and Ca_V_1.2) (Brown and Difrancesco, 1980; DiFrancesco, 1986; Huser et al., 2000; Mangoni et al., 2006), and a “Ca^2+^ clock,” timed by stochastic, diastolic Ca^2+^ sparks from the sarcoplasmic reticulum (SR) that activate inward Na^+^/Ca^2+^-exchanger current (Bogdanov et al., 2001). These coupled oscillators depolarize the membrane to threshold, trigger a cell-wide, global intracellular Ca^2+^ ([Ca^2+^]_i_) transient and, after repolarization by K^+^ currents, reset to begin the next diastolic depolarization.

A defining feature of the SA node is its functional and anatomical heterogeneity: inferior regions fire more slowly and are more sparsely vascularized. This spatial organization creates subpopulations with distinct intrinsic rates, metabolic throughput, and coupling, suggesting that network rhythm could arise through at least two non-exclusive mechanisms: classical entrainment (Jalife, 1984; Anumonwo et al., 1991)—where electrotonic coupling pulls disparate oscillators toward a common period—and stochastic resonance—where variability from a noisier subpopulation can enhance sensitivity and bandwidth (Clancy and Santana, 2020; Grainger et al., 2021; Guarina et al., 2022).

Each beat carries a steep energetic cost. Adenosine triphosphate (ATP) hydrolysis is required to maintain Na^+^ and K^+^ gradients via the Na^+^/K^+^ ATPase, refill the SR with Ca^2+^ through the sarco/endoplasmic reticulum Ca^2+^ ATPase (SERCA) pump, support cross-bridge cycling, and fuel the continual adenylate cyclase-mediated production of cAMP that tunes both clock components (Yaniv et al., 2013). Even under basal conditions, isolated mouse SA node myocytes consume more O_2_ than ventricular myocytes paced at 3 Hz, and their ATP turnover rises further during sympathetic stimulation (Yaniv et al., 2013). About 95% of this ATP is supplied by mitochondrial oxidative metabolism and the remaining ∼5% by glycolysis (Wisneski et al., 1990; Saddik and Lopaschuk, 1991; Zhou and Tian, 2018; Lopaschuk et al., 2021).

Classic work has established coupling between Ca^2+^ handling and ATP production, showing that Ca^2+^ entering the mitochondrial matrix via the mitochondrial Ca^2+^ uniporter (MCU) activates three Ca^2+^-sensitive dehydrogenases—pyruvate, isocitrate, and 2-oxoglutarate dehydrogenase—boosting NADH production and driving proton flux through the ATP5A-containing F_1_F_o_-ATP synthase (Denton et al., 1980; Denton and McCormack, 1990; McCormack et al., 1990; Kirichok et al., 2004; Garbincius and Elrod, 2022). In this process, mitochondrial efficiency is shaped not only by enzymatic capacity but also by organelle architecture and proximity to SR Ca^2+^-release sites. The tethering protein mitofusin-2 (Mfn2), located on the outer mitochondrial membrane, mediates physical and functional coupling between mitochondria and the junctional SR, facilitating rapid Ca^2+^ transfer and enhancing oxidative phosphorylation (Chen et al., 2012; Dorn et al., 2015). Mitochondrial ATP production is supplemented by glycolysis and is further supported by creatine and adenylate kinase reactions, which draw on phosphocreatine and ADP pools to regenerate ATP, with adenine nucleotide translocase (ANT) coupling these processes by exporting mitochondrial ATP in exchange for cytosolic ADP. These pathways act in concert to maintain ATP supply across varying workloads.

To match the relentless demand for ATP by SA node myocytes, the SA node artery (most commonly a branch of the right coronary artery) fans into an elaborate capillary network that perfuses the node during diastole when extravascular compression is lowest. High-resolution, cleared-tissue reconstructions show that capillary density is greatest and myocyte-to-vessel distance shortest in the superior SA node, whereas inferior regions are more sparsely vascularized (Grainger et al., 2021; Manning et al., 2025). These structural gradients correlate with intrinsic firing rate, suggesting that local energetic support sculpts pacemaker hierarchy. Yet direct links between microvascular architecture, subcellular energetics, and pacemaking behavior have remained elusive.

Using genetically encoded fluorescent ATP sensors, we previously showed that, under physiological workload, cytosolic ATP rises and falls with every [Ca^2+^]ᵢ transient in ventricular myocytes (Rhana et al., 2024; Rhana et al., 2025). This raised two questions for the SA node: Do spontaneously firing myocytes exhibit comparable beat-locked ATP dynamics? And do these dynamics differ between superior and inferior domains in ways that would be expected to bias rate setting versus bandwidth?

To address these questions, we combined beat-resolved ATP imaging in the intact node and in isolated myocytes to (i) define two mitochondrial ATP modes time-locked to Ca^2+^ transients; (ii) identify matched high- and low-gain cytosolic ATP phenotypes; (iii) map these energetic phenotypes onto regional firing rate, mitochondrial volume, and capillary density; and (iv) demonstrate that Ca^2+^ release—via SR-to-mitochondria Ca^2+^ transfer—acts as a master regulator that times oxidative ATP production to demand. This hierarchy allocates pacemaking roles across the node and extends the emerging “paycheck-to-paycheck” model from ventricular myocytes (Rhana et al., 2024; Rhana et al., 2025; Santana and Earley, In press) to the SA node. Here, “paycheck-to-paycheck” denotes beat-locked ATP synthesis/use bounded by a local O_2_ ceiling—capillaries set the ceiling, mitochondria the bandwidth, and Ca^2+^ the timing—linking vascular architecture and mitochondrial organization to excitability and clarifying why some cells set rate while others widen bandwidth.

## Methods

### Animals

Male wildtype C57BL/6J mice (The Jackson Laboratory, USA) aged 8–12 weeks were used for all experiments in this study. Mice were euthanized with a single, intraperitoneal administration of a lethal dose of sodium pentobarbital (250 mg/kg). All procedures were performed in accordance with NIH guidelines and were approved by the Institutional Animal Care and Use Committee of the University of California, Davis.

### AAV-mediated delivery of ATP and voltage biosensors

Real-time visualization of intracellular ATP dynamics was enabled by systemically injecting mice with adeno-associated virus serotype 9 (AAV9) vectors encoding either the cytosolic ATP sensor iATPSnFR1.0 (cyto-iATP) (Lobas et al., 2019), the mitochondria-targeted variant iATPSnFR2.0 (mito-iATP) (Marvin et al., 2024), or the ultra-fast, sensitive voltage sensor ASAP5 (Hao et al., 2024) at a titer of 4 x 10^12^ viral genomes per milliliter (vg/mL). AAV9 vector solution (100 μL) was delivered under isoflurane anesthesia (5% induction, 2% maintenance) via retro-orbital injection using a microsyringe. Importantly, iATPSnFR1.0 and iATPSnFR2.0 sensors bind ATP with high specificity and exhibit negligible sensitivity to physiological fluctuations in free [Mg^2+^] (0–5 mM) (Lobas et al., 2019; Marvin et al., 2024). This insensitivity ensures that the observed fluorescence transients reflect genuine changes in ATP availability rather than artifacts arising from concurrent Mg^2+^ fluxes associated with ATP hydrolysis.

### Multi-photon imaging of [ATP]_i_, [ATP]_mito_, and FAD

Following confirmation of deep anesthesia, beating hearts were excised from animals expressing AAV9 constructs and transferred to warmed (37°C) Tyrode III solution consisting of 140 mM NaCl, 5.4 mM KCl, 1 mM MgCl_2_, 1.8 mM CaCl_2_, 5 mM HEPES, and 5.5 mM glucose (pH 7.4, adjusted with NaOH). The SA node was dissected from the beating heart, pinned flat to expose the full node, and incubated in 10 μM blebbistatin (Sigma Aldrich, USA) for 1 hour at 37°C in a humidified 5% CO_2_ atmosphere. Cyto-iATP and mito-iATP signals in intact SA nodes perfused *ex vivo* with room temperature Tyrode III supplemented with blebbistatin (10 μM) were imaged in line-scan mode using an Olympus multi-photon excitation FVMPE-RS upright microscope equipped with an Olympus XL Plan N 25X lens (NA = 1.05). For experiments employing ivabradine (30 μM; Tocris, USA) or thapsigargin (1 μM; Tocris, USA), agents were added directly to the perfusate, and preps were allowed to incubate for 30 minutes before line scans were recorded.

For motion artifact controls, isolated cells and intact SA node preparations co-expressing mito-iATP and HaloTag were imaged using 488-nm (mito-iATP) and 561-nm (HaloTag) excitation after rapid excision and transfer to warm (37 °C) Tyrode’s solution. The ATP sensor was selectively labeled by incubating cells and whole SA nodes for 10 minutes with the JFX554 HaloTag ligand (#HT1030) (Promega, USA) at a final concentration of 0.1 μM. The SA node was carefully dissected free and pinned flat in a recording chamber. Contractile motion was suppressed throughout imaging by maintaining tissues in Tyrode solution supplemented with blebbistatin (10 µM). FAD autofluorescence was acquired using an Olympus FVMPE two-photon microscope equipped with 4x XLFLUOR and 25x XLPLN WMP2 objectives. Two-photon excitation was provided at 910 nm, and emitted fluorescence was collected using non-descanned detectors. Emission was spectrally separated using a 570 nm dichroic mirror (SDM570), and FAD autofluorescence was collected through a 495–540 nm band-pass filter (BA495–540). Images were acquired at a resolution of 512 x 512 pixels. SA nodes were continuously perfused with warmed Tyrode solution containing blebbistatin during imaging. Z-stacks were collected with a 2-µm step size from both the superior and inferior regions of the SA node. For analysis, maximum-intensity z-projections were generated from each z-stack. Regions of interest were defined within the SA node, and mean FAD autofluorescence intensity was quantified after background subtraction.

Fluorescence values were normalized to the pre-treatment control image (F/F_0_). In a subset of experiments, cyto-iATP fluorescence signals were converted to concentration units using the “F_max_” equation (Maravall et al., 2000):

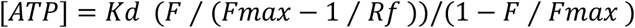

where F is fluorescence, F_max_ is the fluorescence intensity of the iATP sensor in the presence of a saturating ATP concentration, K_d_ is the apparent dissociation constant of the fluorescence indicator used, and R_f_ is the indicator’s dynamic range. In the present study, F_max_ was determined under nodal imaging conditions by superfusing β-escin–permeabilized SA node preparations with 10 mM ATP. The K_d_ (1460 μM) and R_f_ (3.8) values used were empirically determined in ventricular myocytes (Rhana et al., 2024).

### Electrical field stimulation and metabolic/voltage imaging of isolated SA node

Electrically evoked activity in isolated SA node preparations was induced by applying field stimulation using two platinum wire electrodes positioned in close proximity on opposite sides of the SA tissue within the perfusion chamber. The SA node was dissected and prepared as described above and continuously superfused with warmed Tyrode solution containing blebbistatin (10 µM) to limit motion. Electrical stimuli were generated by delivering voltage pulses with a duration of 6 ms using a Grass stimulator (AstroMed Inc, USA). Baseline measurements of cytosolic ATP (cyto-iATP) and membrane voltage (ASAP5) were first acquired under sinus rhythm. The preparation was then entrained by applying field stimulation at 3 Hz, and cyto-iATP and ASAP5 signals were recorded during steady-state pacing. Following baseline and paced recordings, the mitochondrial uncoupler FCCP was added to the perfusate. During FCCP exposure, the effects of metabolic stress and increasing electrical demand on cyto-iATP and membrane voltage dynamics were assessed by incrementally increasing stimulation voltage to 10, 30, and 70 V. Stimulation intensity at each voltage was maintained only for the duration required to obtain stable cyto-iATP line-scan recordings in the SA node.

### Whole-mount immunolabeling of SA node tissue

Immunohistochemistry analyses of the SA node were performed as described previously in detail (Grainger et al., 2021; Manning et al., 2025). Briefly, the SA node tissue was fixed in 4% paraformaldehyde in phosphate-buffered saline (PBS) for 30 minutes. After fixation, the tissue was washed three times for 5 minutes each and then transferred to a 15-mL tube filled with PBS and washed for 12 hours at low speed on a tube rotator. Tissues were dehydrated through a graded ethanol series (25%, 50%, 75%, 95%, 100%), cleared in 20% dimethyl sulfoxide (DMSO) in ethanol for 2 hours, and bleached overnight (12 hours) in 6% hydrogen peroxide prepared in absolute ethanol. Samples were then rehydrated by reversing the ethanol gradient and washed in PBS (3 x 5 minutes). Cells were permeabilized using 0.5% Triton X-100 in PBS (3 x 10 minutes) and then blocked by incubating with 5% normal donkey serum in PBS containing 0.1% Triton X-100 for 2 hours at room temperature.

For immunolabeling, SA nodes were incubated at 4°C for 48 hours with goat anti-mouse CD31 primary antibody (AF3628; R&D Systems, USA), diluted 1:50 in PBS. After three PBS washes (10 minutes each), tissues were incubated for 4 hours at room temperature in the dark with donkey anti-goat Alexa Fluor 568 (1:1000, A11057; Thermo Fisher Scientific, USA) or anti-GFP Alexa Fluor 647 (1:1000, A32985; Thermo Fisher Scientific, USA) secondary antibodies. Following final washes in PBS (3 x 10 minutes), tissues were incubated in a 1:4 solution of DMSO:PBS for 2 hours, then mounted in Aqua-Mount (Thermo Fisher Scientific, USA), cover-slipped, and sealed for imaging.

### Isolation of SA node myocytes

The full SA node was dissected from the beating heart and exposed by pinning flat. The procedure for isolating single SA node myocytes was adapted from Fenske et al. (2020) and Grainger et al. (2021), with modifications. The node was incubated at 37°C for 5 minutes in low-Ca^2+^ Tyrode solution consisting of 140.0 mM NaCl, 5.4 mM KCl, 0.5 mM MgCl₂, 0.2 mM CaCl_2_, 5.0 mM HEPES, 5.5 glucose, 1.2 mM KH_2_PO_4_, and 50.0 mM taurine (pH 6.9, adjusted with NaOH). Enzymatic digestion was performed for 25 minutes at 37°C with periodic (every 5 minutes) gentle agitation using the same Ca^2+^-free Tyrode buffer (pH 6.9) supplemented with 18.87 U/mL elastase, 1.79 U/mL protease, 0.54 U/mL collagenase B, and 100 mg/mL bovine serum albumin (BSA). Following enzymatic treatment, tissues were rinsed twice in fresh low-Ca²⁺ Tyrode and twice in ice-cold Kraft-Brühe (KB) solution consisting of 100.0 mM L-glutamic acid (K^+^ salt), 5.0 mM HEPES, 20.0 mM glucose, 25.0 mM KCl, 10.0 mM L-aspartic acid (K^+^ salt), 2.0 mM MgSO_4_, 10 mM KH_2_PO_4_, 20 mM taurine, 5.0 mM creatine, 10.0 mM EGTA, and 1.0 mg/mL BSA (pH 7.4, adjusted with KOH). Samples were stored at 4°C in 350 μL KB solution for 2–3 hours, then warmed to 37°C for 10 minutes prior to gentle mechanical dissociation using a fire-polished glass pipette. Ca^2+^ was reintroduced in two steps performed at room temperature. First, 10 μL of Na^+^/Ca^2+^ solution containing 10.0 mM NaCl and 1.8 mM CaCl_2_ was added for 5 minutes, then 23 μL of Na^+^/Ca^2+^ solution was added and incubated for 5 minutes. A BSA-containing solution consisting of 140.0 mM NaCl, 5.4 mM KCl, 1.2 mM KH_2_PO_4_, 1.0 MgCl₂, 1.8 mM CaCl_2_, 5.0 mM HEPES, 5.5 mM glucose, and 1 mg/mL BSA was added at volumes of 55 µL, 175 µL and 612 µL in 4-minute steps.

### Confocal imaging of [Ca^2+^]_i_, [ATP]_i_, [ATP]_mito_, and mitochondria

Diastolic and systolic [Ca^2+^]_i_ and cyto-iATP or mito-iATP signals from dissociated SA node myocytes were simultaneously recorded using an inverted Olympus FV3000 confocal microscope (Olympus, Japan) operating in 2D or line-scan mode. Cytosolic Ca^2+^ was monitored by first loading SA node myocytes expressing cyto-iATP or mito-iATP with 3 µM Rhod-3-AM (Thermo Fisher Scientific, USA), as described previously (Rhana et al., 2024). After loading cells with Rhod-3, a drop of the cell suspension was transferred to a temperature-controlled (37°C) perfusion chamber on a microscope stage, and cells were allowed to settle onto the coverslip for 5 minutes. Fluorescent indicators were excited with solid-state lasers emitting at 488 nm (cyto-iATP and mito-iATP) or 561 nm (Rhod-3) through a 60× oil-immersion lens (PlanApo; NA = 1.40). During analysis, background was subtracted from all confocal images, and fluorescence signals were expressed as F/F_0_, where F is the fluorescence intensity at a given time point and F_0_ is the mean baseline fluorescence. Ca^2+^, cyto-iATP, and mito-iATP signals were automatically detected and analyzed using ImageJ and SanPy software (Guarina et al., 2024) or custom Python code.

In a subset of experiments, Ca^2+^ release events were recorded from SA node myocytes loaded with Fluo-4-AM. These signals were detected and analyzed using an ML-based Ca^2+^ signal detection and analysis pipeline as previously described (Garrud et al., 2024). In brief, a F/F_0_ channel was calculated from the raw 488 channel by dividing each frame by a five-frame moving average (without background subtraction) and rescaled to eliminate fractional pixel values. Both channels (488 and F/F_0_) were used to simultaneously train a ML model ground truth by manually annotating every pixel of at least 10 Ca^2+^ signals from 10 representative experiments (Arivis Vision4D, Zeiss). The trained ML model was run on all experiments, and Ca^2+^ signals with a mean F/F_0_ intensity of less than 1.025 of baseline or a total area of less than 5 microns were excluded from further analysis. ML-annotated Ca^2+^ signal properties (size, location [x, y, t], duration, and signal mass [sum of each Ca^2+^ signal F/F_0_ over its duration]) were exported for summary and statistical analysis.

### Super-resolution imaging of SA node myocytes

Freshly isolated SA node myocytes expressing cyto-iATP or mito-iATP were incubated with 250 nM MitoTracker Deep Red (Thermo Fisher Scientific, USA) for 30 minutes at 37°C. Super-resolution radial fluctuation (SRRF) imaging was performed using a Dragonfly 200 spinning-disk confocal system (Andor Technology, UK) equipped with an Andor iXon EMCCD camera. This system was coupled to an inverted Leica DMi microscope equipped with a 60x oil-immersion objective with a numerical aperture (NA) of 1.4. Images were acquired using Fusion software (Andor, UK). Colocalization of ATP sensors with MitoTracker was evaluated using the colocalization module in Imaris 10 (Oxford Instruments).

### Quantification of mitochondrial volume

Mitochondrial volume was quantified by analyzing SRRF-acquired 3D image stacks of MitoTracker-labeled SA node myocytes with Imaris 10 software (Oxford Instruments, UK). Myocytes were isolated from anatomically defined superior and inferior regions of the SA node. Mitochondria were segmented using the Surfaces tool based on MitoTracker Deep Red fluorescence intensity, applying a consistent threshold across all samples. Total mitochondrial volume was computed for each individual cell, allowing for regional comparisons between superior and inferior SA node myocyte populations.

### SA node cryosectioning

The SA node region was fixed in 4% paraformaldehyde for 30 minutes, washed in PBS (0.01 M), and dehydrated in PBS containing 30% sucrose for 24 hours prior to embedding in OCT embedding medium (Sakura, USA). Tissue was oriented perpendicular to the long axis of the SA node during embedding for longitudinal sectioning. Cryosections were cut at 5 μm thickness and mounted onto glass slides for subsequent *in situ* hybridization.

### RNAscope in situ hybridization

*In situ* hybridization was performed using the RNAscope 2.5 HD Duplex Assay Kit (Advanced Cell Diagnostics, USA) with the Mm-Atp5a1-C2 probe (#459311-C2) and hematoxylin counterstaining, following the manufacturer’s standard protocol. Atp5a1-positive puncta were visualized using Fast Red chromogenic detection and their fluorescence was imaged on an APX100 microscope equipped with a UPLXAPO 40x objective (NA 0.95; Olympus, Japan) using a TRITC filter cube. A single projected image resolving puncta throughout the tissue thickness was generated from Z-stacks spanning the full section thickness (5 μm, 14 optical planes), acquired using extended focal imaging and stitching functions in CellSens APEX 4.3 software. Image analysis was performed using ImageJ version 2.14.0 (NIH, USA). Rectangular regions of interest (ROIs; 250 x 25 μm) were positioned along the long axis of the SA node. ROIs were pre-processed using a 2-pixel radius median filter followed by rolling-ball background subtraction (radius, 30 pixels). Local fluorescence maxima, identified using the Find Maxima function with a fixed prominence threshold, were used to detect both isolated and clustered Atp5a1 RNA puncta. The total number of detected maxima within each ROI was normalized to ROI area and reported as puncta density (puncta per mm^2^). Hematoxylin counterstaining was used for anatomical reference and confirmation of SA node regional identity.

### Statistics

Data were analyzed using hierarchical statistics to account for the nested structure of the data (cells within animals). Per-cell values were first summarized within each animal, and animals were treated as the unit of inference (*N* in figures represents the number of animals). Where variance differed between groups (heteroscedasticity assessed by Brown–Forsythe), Welch’s ANOVA or Welch’s t-test with Games–Howell *post hoc* comparisons were used. Distributions were analyzed using finite Gaussian-mixture models, with component selection based on the Bayesian information criterion. Statistical significance was set at *P* < 0.05; individual P-values are denoted in figures. Distributions that were statistically multimodal (i.e., ATP signal mass, Ca^2+^ signal mass, and mitochondrial volume fraction) were modelled with finite Gaussian-mixture models. The optimal number of components was selected by minimizing the Bayesian information criterion and confirmed by the gap statistic, silhouette score, and Hartigan’s dip test for unimodality (*P* < 0.001 for departure from a single mode).

### Online supplemental material

Online Supplemental **Figures S1** and **S2** quantify the kinetics of beat-locked ATP signals in the intact SA node, with **Figure S1** reporting the rise/decay parameters of whole-node cyto-iATP transients and **Figure S2** reporting the corresponding kinetic analysis of whole-node mito-iATP Mode 1 and Mode 2 events. Supplemental **Figure S3** provides ratiometric controls using ATP-insensitive HaloTag to validate that beat-locked mito-iATP oscillations are not driven by motion or optical-path fluctuations in intact nodes. Supplemental

Figures S4 and S5 focus on acutely dissociated SA node myocytes and relate intracellular Ca^2+^ handling to ATP dynamics at the single-cell level. Supplemental **Figure S6** provides ratiometric controls using ATP-insensitive HaloTag to validate that beat-locked mito-iATP oscillations are not driven by motion or optical-path fluctuations in isolated myocytes. Finally, **Figure S7** shows the kinetic analysis of Ca^2+^ and mito-iATP signals in high- and low-load sites.

## Results

### Superior and inferior regions of the SA node exhibit distinct cytosolic ATP dynamics

To map regional ATP dynamics, we first expressed the genetically encoded cytosolic ATP reporter, iATPSnFR1 (cyto-iATP) (Lobas et al., 2019), in mouse SA node myocytes, then surveyed cyto-iATP fluorescence throughout intact mouse SA node whole mounts (Figure 1). Preparations were co-stained with the endothelial marker, CD31, to provide vascular landmarks and demarcate SA node borders. In agreement with prior work (Grainger et al., 2021; Manning et al., 2025), vessel density varied markedly along the mouse node and was highest in the superior region (Figure 1A). Cyto-iATP signals were detected throughout the node, and cyto-iATP fluorescence was diffusely distributed in isolated SA node myocytes. Automated segmentation revealed more cyto-iATP–positive myocytes in the superior than the inferior region, consistent with the previously reported SA node myocyte density gradient; however, mean cyto-iATP fluorescence per cell did not differ between superior and inferior regions (Figure 1B**, C**).

**Figure 1.**
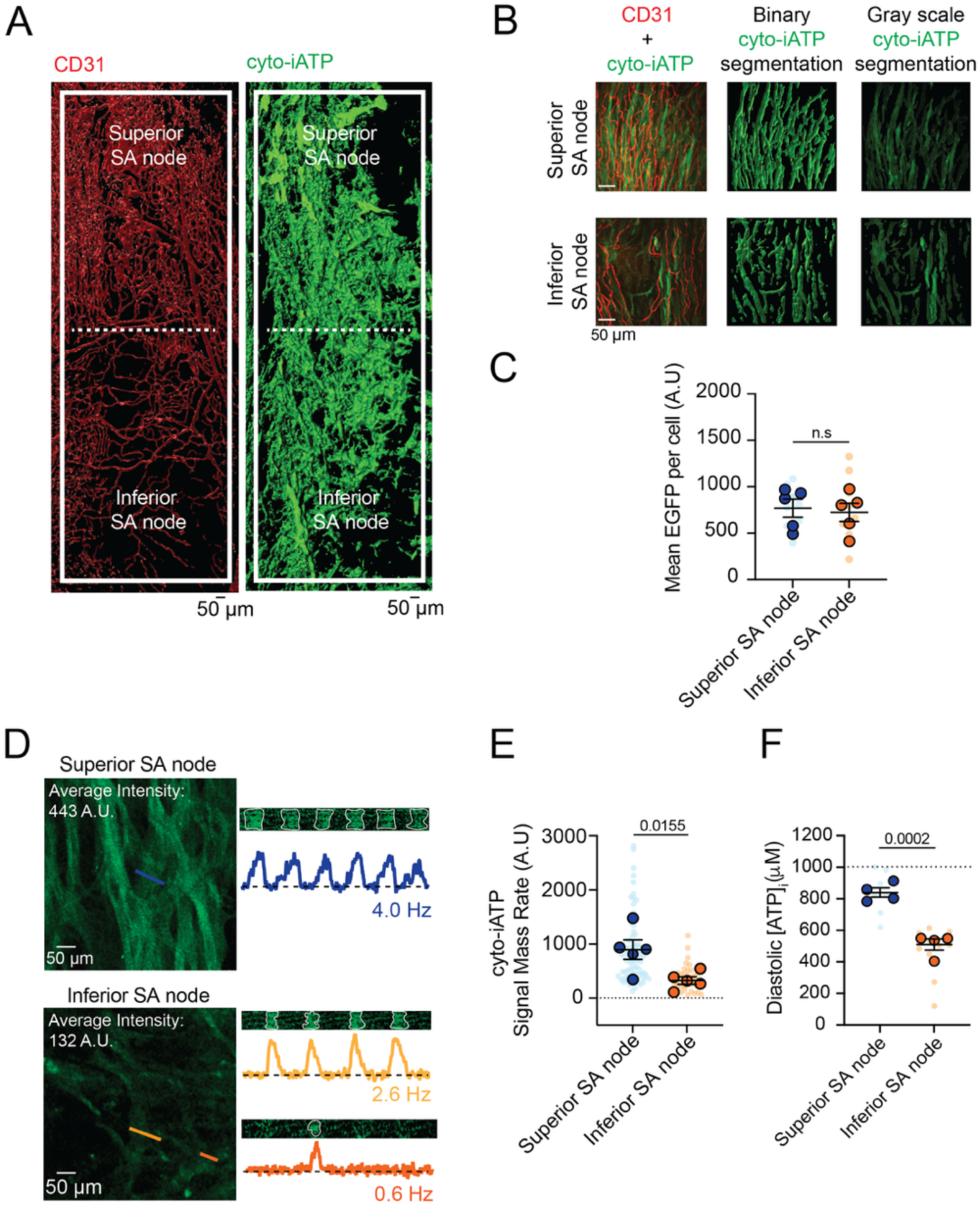
Superior SA node myocytes exhibit elevated diastolic ATP and metabolic flux compared with the inferior region. (**A**) 3D segmented maximum-intensity projection of a whole-mount SA node immunolabeled for CD31 (vasculature, red) and cyto-iATP (myocytes, green). The dashed line denotes the boundary between superior and inferior regions. (**B**) Image-processing workflow illustrating merged maximum-intensity projections, binary segmentation masks, and extraction of grayscale cyto-iATP signals used for quantitative analysis. (**C**) Mean cyto-iATP fluorescence intensity per myocyte, grouped by region (N = 5 mice per region), reporting expression levels of the EGFP-tagged cyto-iATP sensor. (**D**) Live confocal imaging of cyto-iATP signals showing representative line-scan images and corresponding normalized fluorescence traces (F/F₀) from superior and inferior regions. (**E**, **F**) Summary quantification of cyto-iATP signal mass rate (**E**) and estimated diastolic [ATP]_i_ (**F**). *P*-values are shown above comparisons. Large circles denote per-animal means; small circles indicate individual biological replicates.

We next imaged cyto-iATP *ex vivo* in intact SA nodes during sinus rhythm using multiphoton microscopy while suppressing contraction with blebbistatin (10 µM) to eliminate motion artifacts. In these live preparations, optical sections consistently exhibited higher regional mean cyto-iATP fluorescence in the superior domain than in the inferior domain (Figure 1D). Because mean per-cell EGFP-tagged cyto-iATP fluorescence did not differ in the whole-mount analysis (Figure 1C), the elevated regional signal in live tissue is consistent with the hypothesis that steady-state cytosolic ATP availability is higher in the superior pacemaker zone.

Line-scan multi-photon imaging further revealed large, rhythmic cyto-iATP transients in the superior SA node that tracked the local firing rate (∼4 Hz; Figure 1D**, top**). Inferior regions were more heterogeneous: some myocytes fired more slowly (∼2.6 Hz), whereas others were nearly quiescent (∼0.6 Hz; Figure 1D**, middle and bottom**). Although action potentials were not recorded in these experiments, the observed frequencies closely match reported electrical measurements in superior and inferior SA node myocytes, supporting the interpretation that cyto-iATP oscillations are beat-locked readouts of intrinsic pacemaker activity (Grainger et al., 2021).

A kinetic analysis showed similar time-to-peak for superior (67.85 ± 2.11 ms) and inferior (70.14 ± 3.05 ms) cyto-iATP transients (*P* = 0.2777; **Figure S1**). In contrast, decay phases, measured as time to 50% amplitude (t_1/2_), were slower in inferior cells (t_1/2_ = 30.25 ± 3.22 ms) than superior cells (t_1/2_ = 20.55 ± 1.52 ms; *P* = 0.0181), yielding a longer overall transient duration in the inferior region (183.70 ± 10.53 ms) compared with the superior region (147.80 ± 6.41 ms; *P* = 0.0120).

To quantify beat-to-beat energetic output independent of absolute calibration, we computed the cyto-iATP signal mass of each transient, defined as the time- and area-integrated change in fluorescence. A compilation of all individual transients over identical recording durations (4.2 seconds), showed that the total cyto-iATP signal mass was markedly greater in cells of the superior SA node (895.70 ± 180.80 AU) than in inferior cells (326.40 ± 71.75 AU). Consistently, across nodes, the superior region exhibited a higher signal-mass rate (total signal mass per trace divided by recording time; Figure 1E; *P* = 0.0155). Thus, superior SA node myocytes sustain higher beat-to-beat energetic throughput than inferior myocytes.

To estimate regional differences in absolute ATP concentration, we converted cyto-iATP fluorescence to [ATP]_i_ using an F_max_-based approach (Maravall et al., 2000). F_max_ was determined by permeabilizing SA nodes expressing cyto-iATP with β-escin followed by exposure to saturating (10 mM) ATP; [ATP]_i_ was then computed by combining F_max_ with prior *in situ* sensor parameters (Rhana et al., 2024). Using this approach, we found that diastolic [ATP]_i_ was sub-millimolar in both regions but was significantly higher in the superior SA node (839.9 ± 29 µM) than in the inferior SA node (510.1 ± 35.24 µM; *P* = 0.0002), consistent with the higher calibration-independent cyto-iATP signal-mass rate in superior cells (Figure 1F).

### Pharmacological dissection of I_f_ and SR Ca^2+^ contributions to cytosolic ATP dynamics

We next asked how two key contributors to pacemaker cycling—If-dependent depolarization and SR Ca^2+^ cycling/whole-cell Ca^2+^ transients shape energetic demand in the SA node (Figure 2). To address this, we imaged cyto-iATP–expressing nodes in sinus rhythm before and after inhibiting I_f_-producing HCN channels with ivabradine or suppressing SR Ca^2+^ release with the SERCA inhibitor thapsigargin (Figure 2A). Under basal conditions, superior SA node regions exhibited rhythmic cyto-iATP transients at 3.27 ± 0.06 Hz (N = 6 SA nodes), whereas inferior regions fired more slowly at 2.12 ± 0.07 Hz (N = 6 SA nodes; *P* = 0.0203), but their amplitude (1.17 ± 0.02 F/F_0_; *P* = 0.9978) was similar to that of superior cells (1.20 ± 0.01 F/F_0_; *P* = 0.7433).

**Figure 2.**
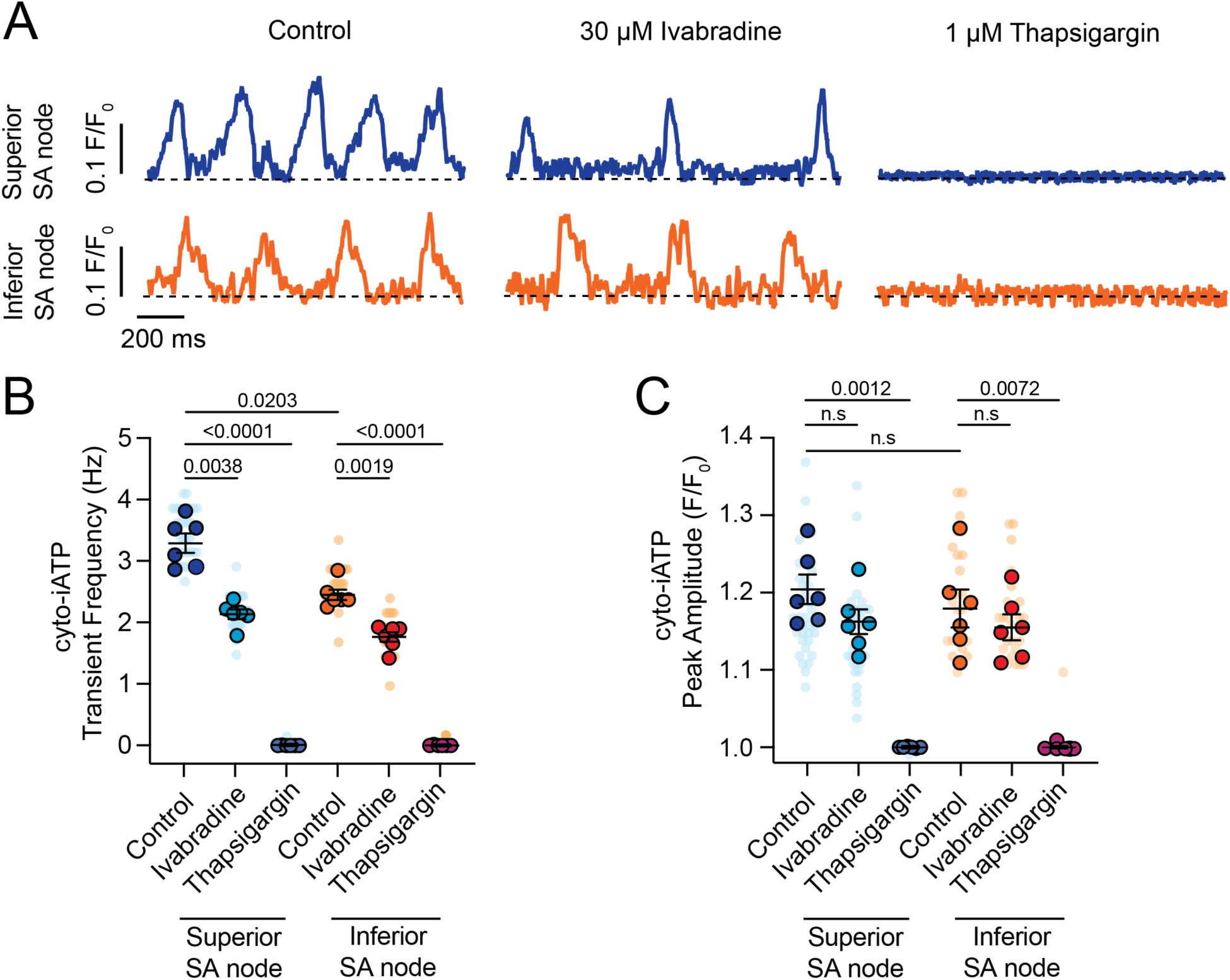
SR Ca^2+^ release drives beat-to-beat ATP oscillations, while I_f_ modulates metabolic frequency. (**A**) Representative normalized cyto-iATP confocal line-scan traces from superior (top) and inferior (bottom) SA nodes acquired under control conditions (left), after I_f_ inhibition with ivabradine (30 µM; middle), and following SERCA inhibition with thapsigargin (1 µM; right). Traces are shown as normalized fluorescence (F/F₀). (**B, C**) Summary quantification of cyto-iATP oscillation frequency (**B**) and peak cyto-iATP oscillation amplitude (**C**) across conditions (N = 6 mice per group). *P*-values are shown above comparisons. Large circles denote per-animal means; small circles indicate individual biological replicates.

Ivabradine (30 µM) lengthened the cycle, decreasing frequency from 3.27 ± 0.06 Hz to 2.12 ± 0.07 Hz in superior cells (*P* = 0.0038) and from 2.45 ± 0.08 Hz to 1.76 ± 0.07 Hz in inferior cells (*P* = 0.0019), but did not alter the transient peak (F/F_0_: 1.16 ± 0.01 and 1.15 ± 0.01 in superior and inferior SA nodes, respectively) (Figure 2B**, C**). Thus, HCN channel inhibition curtailed ATP demand chiefly by lowering beat rate. In contrast, blocking SERCA with thapsigargin (1 µM) abolished spontaneous firing, collapsing cyto-iATP transients to baseline, and prevented recovery during the 10-minute recording (Figure 2B**, C**). The complete loss of ATP pulses despite an intact membrane clock indicates that SR Ca^2+^ cycling is essential not only for excitability but also for beat-to-beat ATP generation.

Together, these manipulations suggest that chronotropic control and SR Ca^2+^ re-uptake make separable, complementary contributions to cellular ATP turnover. The high metabolic output characteristic of the superior pacemaker zone is critically dependent on continuous SERCA activity, underscoring the role of Ca^2+^ release as a master regulator of metabolic supply rather than a passive follower of electrical activity.

### Ca^2+^ transient-locked mitochondrial ATP transients are stronger in superior SA node myocytes

To test whether cyto-iATP oscillations arise from beat-locked changes in mitochondrial ATP, we recorded line-scan images across an intact, mito-iATP–expressing SA node (Figure 3). The mitochondrial reporter, mito-iATP, derived from iATPSnFR2 bearing the A95A/A119L low-affinity (apparent K_d_ = 0.5 mM) variant, was targeted to the matrix by four tandem COX8 leader sequences (Marvin et al., 2024). Figure 3A shows images of an isolated SA node myocyte expressing mito-iATPSnFR2 and stained with MitoTracker. Note that mito-iATPSnFR2 displayed a punctate fluorescence pattern characteristic of mitochondrial structures and strong spatial overlap with MitoTracker Far Red, with a mean Manders’ correlation coefficient of 0.85 ± 0.03, consistent with robust mitochondrial targeting.

**Figure 3.**
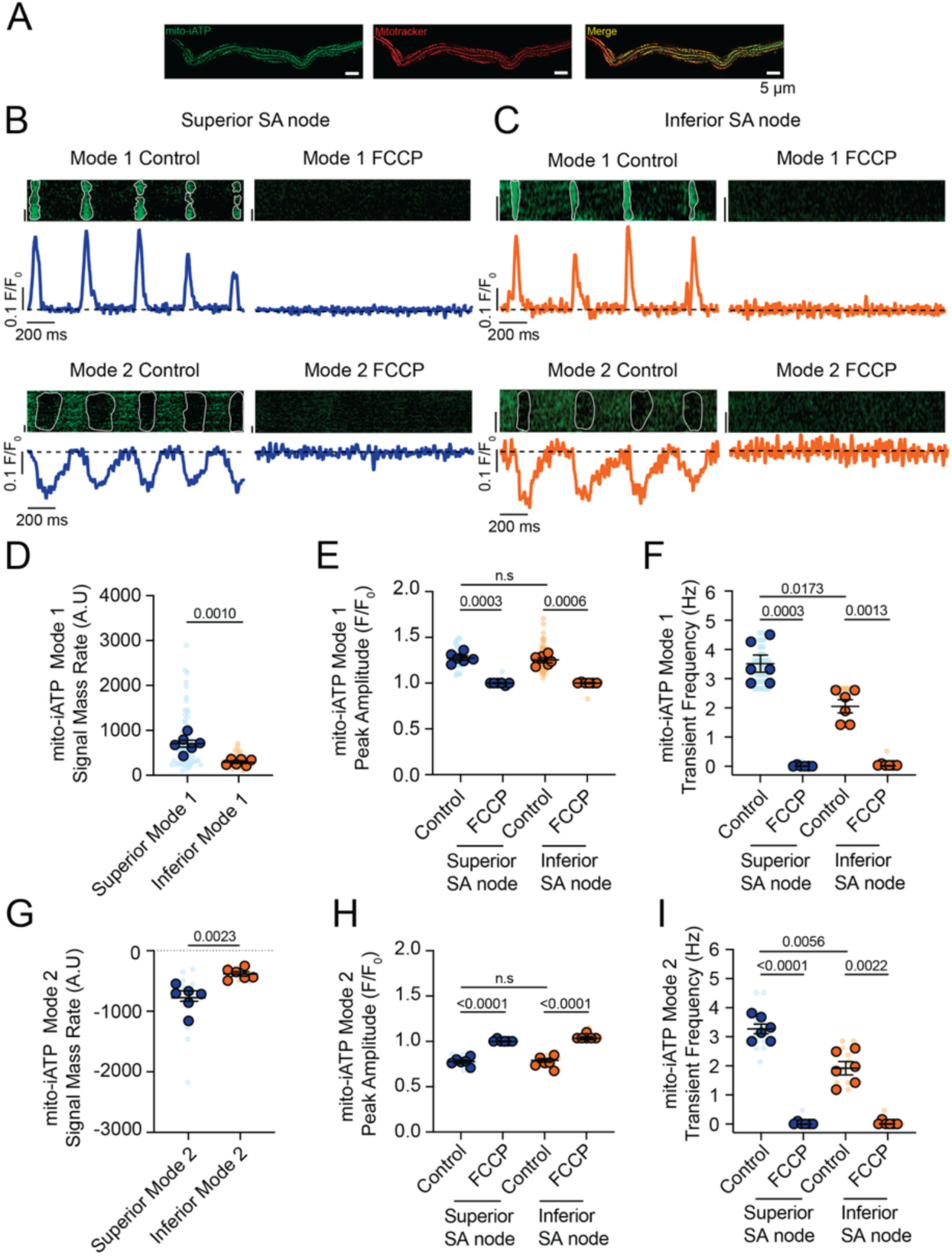
Oxidative phosphorylation drives distinct modes of beat-to-beat ATP dynamics in SA node mitochondria. (**A**) Representative confocal image of an isolated SA node myocyte co-expressing mito-iATP (green) and labeled with MitoTracker (red), illustrating mitochondrial localization of the mito-ATP sensor. Scale bar, 5 µm. (**B**, **C**) Representative confocal line-scan images and corresponding normalized fluorescence traces (F/F₀) from superior (**B**) and inferior (**C**) SA node regions. Two classes of mito-iATP signals are resolved: Mode 1 transients, characterized by positive deflections, and Mode 2 transients, characterized by negative deflections. Traces acquired after application of the mitochondrial uncoupler FCCP (1 µM) are shown for comparison. (**D**–**F**) Summary quantification of Mode 1 mito-iATP dynamics, including signal mass rate (**D**), peak amplitude (**E**), and transient frequency (**F**). (**G**–**I**) Summary quantification of Mode 2 mito-iATP dynamics, including signal mass rate (**G**), peak amplitude (**H**), and transient frequency (**I**). *P*-values are shown above comparisons. Data are presented as means ± SEM (N = 6 mice per group). Large circles denote per-animal means; small circles indicate individual biological replicates.

Having validated the selective mitochondrial targeting of mito-iATP in SA node pacemaker cells, we proceeded to perform high-resolution imaging of mitochondrial ATP dynamics. Spontaneous, rhythmic mitochondrial signals were detected in both superior and inferior regions of the node, appearing as two stereotyped waveforms, designated Mode 1 and Mode 2 (Figure 3B**, C**). Mode 1 events exhibited a rapid rise after each beat, followed by decay, whereas Mode 2 events showed a transient dip before recovering, presumably as new ATP molecules were synthesized.

We performed a detailed kinetic analysis of mito-iATP events in the intact SA node (**Figure S2**). For Mode 1 “gain” events, rise and decay kinetics were statistically indistinguishable in superior and inferior SA nodes; time-to-peak was 69.23 ± 2.29 ms in the superior region and 70.58 ± 0.52 ms in the inferior region, followed by decay with a time to 50% amplitude (t_1/2_) of 22.48 ± 0.86 ms in the superior region and 21.21 ± 1.05 ms in the inferior region (**Figure S2A–E**). Mode 2 “dip” events reached a similar nadir in the superior node (113.70 ± 1.94 ms) and the inferior node (112.30 ± 4.59 ms) (*P* = 0.7784); recovery proceeded with higher t_1/2_ values of 38.49 ± 1.22 ms in the superior region compared to 31.91 ± 1.58 ms in the inferior region (**Figure S2F–J**). The full duration of Mode 1 mito-iATP transients in the superior region was 157.5 ± 2.7 ms, similar to that in the inferior region (155.40 ± 4.13 ms). Mode 2 mito-iATP durations were longer in the superior region (282.20 ± 16.74 ms) than in the inferior region (240.0 ± 8.05 ms; *P* = 0.046). Given that the recovery phase of Mode 2 events reflects oxidative-phosphorylation capacity, the observed ∼240–282 ms recovery window suggests that dip-dominant cells can sustain firing only up to ∼3.2–4.0 Hz, as faster pacing would limit ATP replenishment before the next beat.

The mito-iATP signal mass rate (i.e., total signal mass per trace ÷ recording time) of Mode 1 sites was significantly larger in superior myocytes (704.0 ± 76.92 AU) than inferior myocytes (292.80 ± 28.17 AU), indicating greater mitochondrial ATP production in the leading pacemaker zone (Figure 3D; *P* = 0.0010). Mode 1 mito-iATP amplitudes in the superior region (1.28 ± 0.02 F/F_0_) were similar to those in the inferior region (1.25 ± 0.02 F/F_0_) (Figure 3E; *P* = 0.98). The frequency of Mode 1 mito-iATP transients in the superior region ranged from 1.2 to 4.5 Hz, with a mean frequency of 3.51 ± 0.28 Hz. In the inferior node, mito-iATP transient frequency never exceeded 2.60 Hz, and exhibited a mean frequency of 2.05 ± 0.22 Hz (Figure 3F; *P* = 0.0173).

The integrated ATP deficit rate (i.e., negative signal mass) during Mode 2 events was significantly larger in superior myocytes (-775.90 ± 86.70 AU) than inferior myocytes (-383.10 ± 37.46 AU) (Figure 3G; *P* = 0.0023), indicating greater mitochondrial ATP consumption in the leading pacemaker zone. Dip amplitudes were virtually identical between regions (0.77 ± 0.01 vs. 0.79 ± 0.02 F/F_0_; Figure 3H; *P* = 0.99). Mean Mode 2 event frequencies in superior and inferior nodes were 3.27± 0.17 Hz and 1.91 ± 0.23 Hz, respectively (Figure 3I; *P* = 0.0056), the latter of which is well below the ∼3.2–4 Hz ceiling predicted from the 250–312 ms recovery window.

Thus, in Mode 2 microdomains, ATP is drawn out of the matrix faster than it is produced during the action potential, generating a brief deficit that is repaid more slowly during diastole. Because Mode 1 (gain) and Mode 2 (dip) regions are spatially distinct, this imbalance implies that high-demand microdomains (Mode 2) temporarily outstrip the surplus generated in high-production zones (Mode 1). Matrix diffusion and a diastolic repayment window, therefore, appear essential for beat-to-beat energetic homeostasis in pacemaker cells.

Brief exposure to FCCP (1 µM), an uncoupler of oxidative phosphorylation, abolished both waveform types in every cell examined (Figure 3B**–I**). Thus, beat-locked mitochondrial ATP transients depend on an intact proton-motive force and, therefore, on ongoing oxidative phosphorylation. The complete suppression of Mode 1 signals—and the greater reduction in Mode 2 amplitude—in superior myocytes underscores their heavier reliance on oxidative phosphorylation and highlights the tight electro-metabolic coupling that supports the dominant pacemaker site. By contrast, the data suggest that inferior mitochondria make only a modest contribution to the nodal cytosolic ATP pool under basal conditions.

Intact-node multiphoton imaging can be confounded by residual micro-motion and scan-registration error even with blebbistatin. Accordingly, to confirm that beat-locked mito-iATP fluorescence changes reflect genuine ATP dynamics rather than motion artifacts, we performed a control experiment using an ATP-insensitive reference reporter in the same optical path (**Figure S3**). To this end, we co-expressed mito-iATP with an inert, ATP-insensitive HaloTag protein labeled with the fluorophore, Janelia Fluor JFX554, a red-shifted HaloTag Ligand, and recorded both signals simultaneously. Beat-locked oscillations were observed exclusively in the iATP channel, whereas the HaloTag signal remained flat under identical imaging conditions. Importantly, both Mode 1 and Mode 2 transients in the intact SA node were preserved by ratiometric normalization (mito-iATP/HaloTag), supporting the conclusion that the observed waveforms are not explained by motion or optical-path fluctuations.

### Uncoupling oxidative phosphorylation suppresses electrical excitability and abolishes beat-locked cyto-iATP signals

To determine whether the loss of beat-locked ATP signals during mitochondrial uncoupling reflects direct disruption of ATP production (rather than an optical artifact) or instead represents a failure of electrical activation, we compared cyto-iATP dynamics with a parallel optical readout of membrane voltage (Figure 4). In spontaneously active SA node myocytes, cyto-iATP exhibited rhythmic transients during sinus rhythm (Figure 4A). Imposing field stimulation (3 Hz, 5 V) entrained cyto-iATP transients, indicating that cytosolic ATP signals can be driven by externally paced excitation (Figure 4B). In the presence of the mitochondrial protonophore FCCP (10 µM), cyto-iATP transients were abolished during 3 Hz stimulation at 5 V and remained absent even when stimulus intensity was increased stepwise (20–60 V) (Figure 4C), indicating that mitochondrial uncoupling prevents electrically evoked ATP responses.

**Figure 4.**
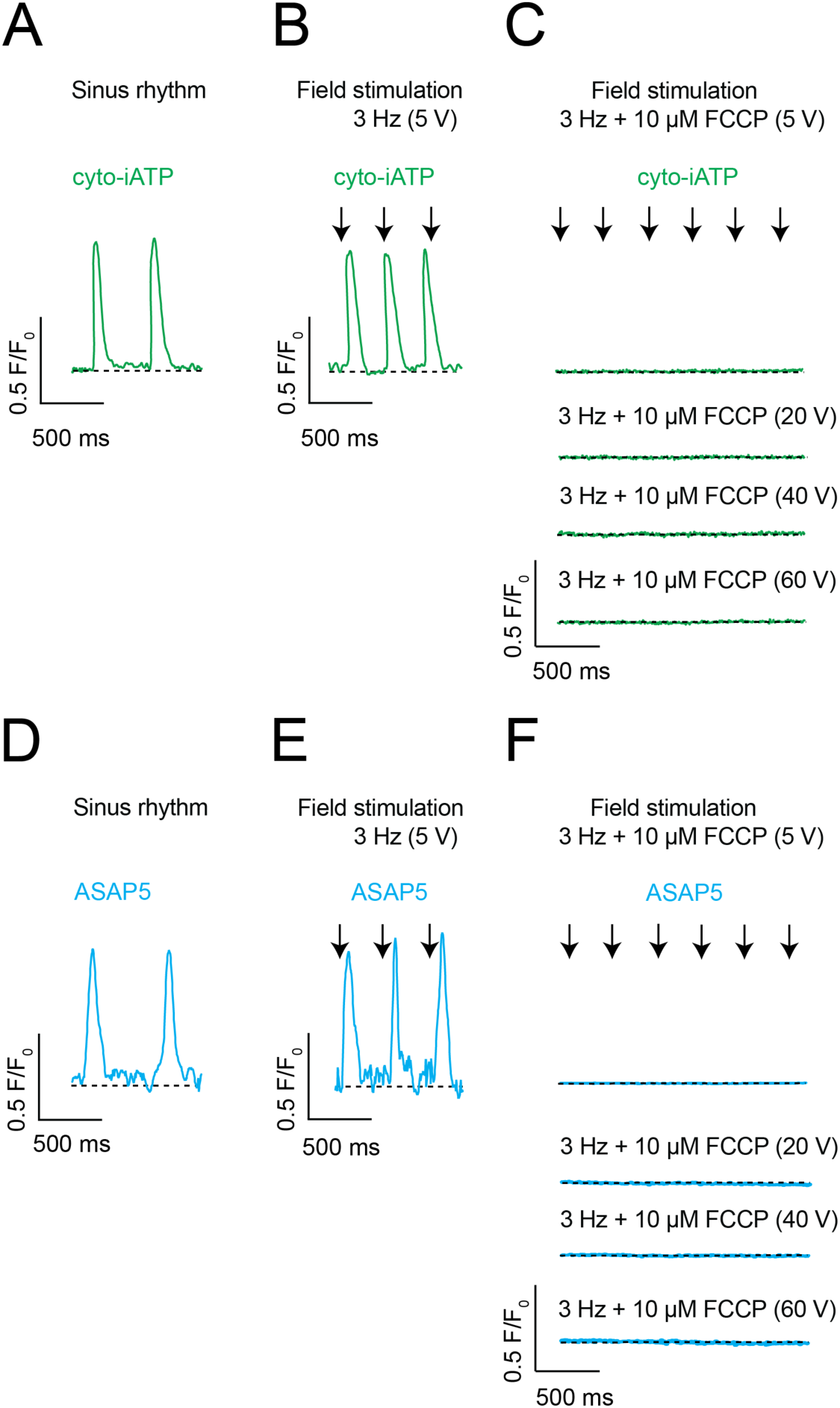
Mitochondrial ATP synthesis is obligatory for SA node excitability and cannot be bypassed by electrical pacing. (**A**–**F**) Representative recordings of cytosolic ATP dynamics (**A**–**C**; cyto-iATP, green) and membrane voltage (**D**–**F**; ASAP5, blue) obtained from intact SA node preparations. (**A**, **D**) Beat-locked cyto-iATP transients with positive deflections and corresponding voltage signals recorded during spontaneous sinus rhythm. (**B**, **E**) Entrainment of cyto-iATP and voltage signals during field stimulation at 3 Hz (5 V). (**C**, **F**) Loss of excitation–metabolism coupling during mitochondrial uncoupling with FCCP (10 µM). Traces were acquired during continued pacing at 3 Hz with stepwise increases in stimulation amplitude (5–60 V). For ASAP5 recordings, upward deflections correspond to membrane depolarization. Traces are shown as normalized fluorescence (F/F₀). Scale bars: 0.5 F/F₀, 500 ms.

Next, using the novel genetically encoded voltage indicator, ASAP5, we tested whether FCCP also disrupts the underlying electrical response to field stimulation (Hao et al., 2024) (Figure 4D**–F**). ASAP5 signals were rhythmic during sinus rhythm and were robustly entrained by 3 Hz field stimulation at 5 V (Figure 4D**, E**). Strikingly, stimulus-locked ASAP5 voltage signals were abolished by FCCP and were not restored by increasing stimulus intensity up to 60 V (Figure 4F). Thus, under our conditions, FCCP renders SA node myocytes electrically unresponsive to field stimulation, providing a parsimonious explanation for the simultaneous loss of beat-locked cyto-iATP signals. Together, these data support the conclusion that intact mitochondrial proton-motive force is required to sustain the excitation–metabolism coupling necessary for beat-locked cytosolic ATP dynamics in pacemaker myocytes.

### Mitochondrial Ca^2+^ uptake and ANT-dependent exchange are required for beat-locked ATP transients

To dissect the downstream sequence linking Ca^2+^ release to ATP production, we targeted the mitochondrial Ca^2+^ uniporter (MCU) and the adenine nucleotide translocase (ANT) (Figure 5). Inhibiting MCU with Ru360 (Matlib et al., 1998; Kim et al., 2011) modestly reduced Ca^2+^ transients (Figure 5A), almost completely abolished cyto-iATP transients (Figure 5B), modestly reduced mitochondrial Ca^2+^ transients (Figure 5C), and abolished Mode-1 mito-iATP gains (Figure 5D). Thus, the gain phase of ATP production is not a passive consequence of increased workload; it requires Ca^2+^ entry through MCU, consistent with Ca^2+^-dependent activation of matrix dehydrogenases and accelerated NADH production on a beat-to-beat timescale. Blocking ANT with bongkrekic acid (BKA) (Henderson and Lardy, 1970; Ruprecht et al., 2019) likewise reduced Ca^2+^ transients (Figure 5A), abolished cyto-iATP transients (Figure 5B), disrupted mitochondrial Ca^2+^ signals (Figure 5C), and abolished mito-iATP transients (Figure 5D), indicating that neither glycolysis nor phosphocreatine buffering can sustain beat-locked ATP oscillations when ADP/ATP exchange across the inner mitochondrial membrane is interrupted. Together, these data demonstrate that a bidirectional MCU–ANT functional unit is essential for both the generation and cytosolic export of beat-locked ATP pulses in the SA node. Together, these data demonstrate that a bidirectional MCU–ANT functional unit is essential for both the generation and cytosolic export of beat-locked ATP pulses in the SA node.

**Figure 5.**
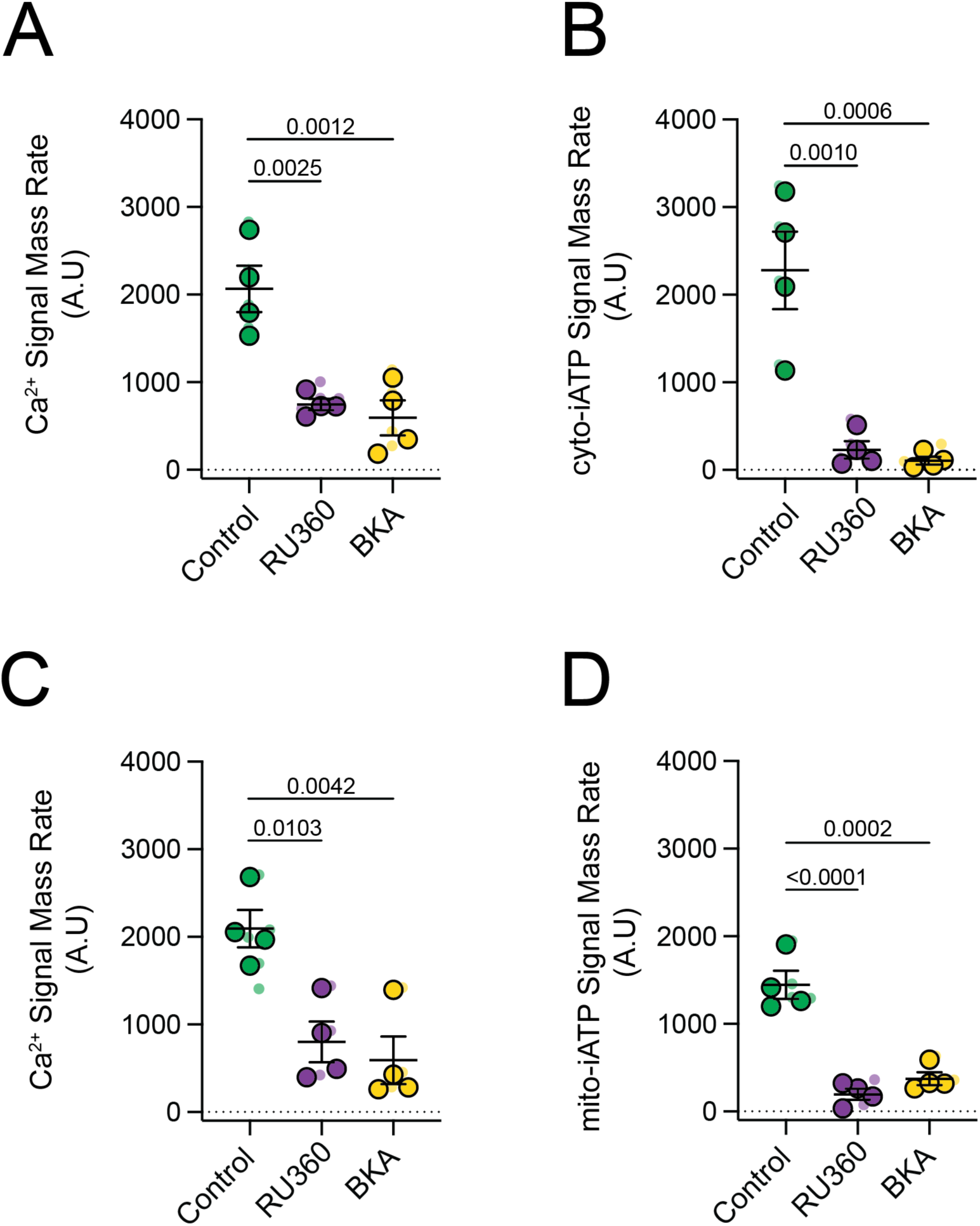
Mitochondrial Ca^2+^ uptake and ANT-mediated ATP export are required for beat-to-beat metabolic signaling. (**A**, **B**) Summary data from SA node myocytes expressing the cytosolic ATP sensor (cyto-iATP). (**A**) Intracellular Ca^2+^ signal mass rates measured under control conditions, after inhibition of the mitochondrial Ca^2+^ uniporter (MCU) with RU360 (5 µM) and following blockade of the adenine nucleotide translocator (ANT) with bongkrekic acid (BKA; 10 µM). (**B**) Corresponding cytosolic ATP signal mass rates obtained under the same conditions. (**C**, **D**) Summary data from parallel experiments in SA node myocytes expressing the mitochondrial ATP sensor (mito-iATP). (**C**) Intracellular Ca^2+^ signal mass rates. (**D**) Mitochondrial ATP signal mass rates. Data are presented as means ± SEM (N = 4 mice per group). *P*-values are shown above comparisons. Large circles denote per-animal means; small circles indicate individual biological replicates.

### Regional coupling of beat-locked ATP dynamics to mitochondrial redox state in the intact SA node

To integrate the superior–inferior gradients in cyto- and mito-iATP with a calibration-independent readout of mitochondrial redox control, we measured endogenous FAD autofluorescence by two-photon microscopy in intact SA node preparations while applying the same perturbations used for ATP imaging (ivabradine, thapsigargin, FCCP) (Figure 6). Regions of interest were defined anatomically as superior and inferior SA node domains (Figure 6A). Because oxidized flavoproteins are fluorescent and these signals are quenched by reduction, increases in FAD autofluorescence report net oxidation of the mitochondrial flavin pool and decreases report net reduction (Huang et al., 2002; Berthiaume et al., 2019).

**Figure 6.**
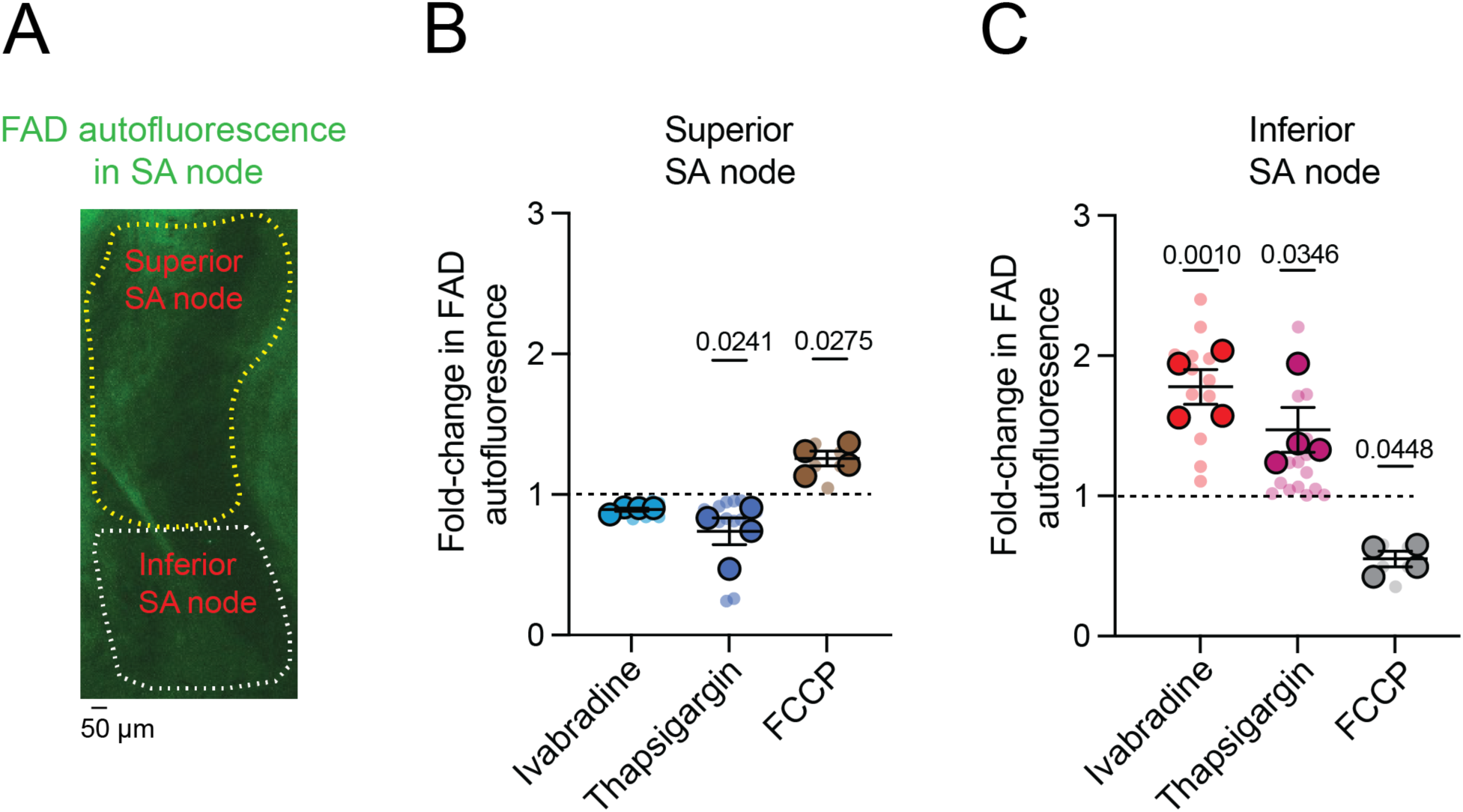
Distinct mitochondrial redox regulation in the superior versus inferior SA node. (**A**) Representative FAD autofluorescence image of a mouse SA node illustrating spatial segregation of the superior (yellow outline) and inferior (white outline) regions. Scale bar, 50 µm. (**B**, **C**) Quantification of normalized FAD autofluorescence (F/F₀) in the superior (**B**) and inferior (**C**) SA node under control conditions and following I_f_ inhibition with ivabradine (30 µM), SERCA inhibition with thapsigargin (1 µM), or mitochondrial uncoupling with FCCP (1 µM). The dashed line indicates baseline fluorescence (F/F₀ = 1). *P*-values are shown above comparisons. *P* values indicate comparisons relative to baseline (F/F₀ = 1). Data are presented as means ± SEM (N = 4 mice). Large circles denote per-animal means; small circles indicate individual biological replicates.

In the superior SA node, where the cyto-iATP signal-mass rate and calibrated diastolic [ATP]_i_ were highest, ivabradine (30 µM) did not significantly alter FAD autofluorescence relative to baseline, whereas thapsigargin (1 µM) produced a significant downward shift, consistent with a net reduction, and FCCP (1 µM) induced the expected oxidizing shift (Figure 6B). Thus, relative to baseline, thapsigargin decreased FAD autofluorescence (*P* = 0.0241), while FCCP significantly increased FAD (*P* = 0.0275).

In contrast, in the inferior SA node, all three interventions significantly altered FAD autofluorescence relative to baseline, both in opposing directions (net oxidation) (Figure 6C). Ivabradine increased FAD autofluorescence (*P* = 0.0010), thapsigargin also increased FAD (*P* = 0.0346), whereas FCCP decreased FAD autofluorescence relative to baseline (*P* = 0.0448).

Together, the region-resolved FAD responses (Figure 6B**, C**) align with the ATP data by indicating that superior and inferior pacemaker zones occupy distinct energetic control regimes: the superior region sustains higher beat-locked ATP throughput and responds to ivabradine or thapsigargin with a net reduction of the flavin pool, consistent with greater energetic reserve, whereas the inferior region exhibits lower ATP availability and shows net flavin oxidation under the same perturbations, consistent with tighter supply–demand constraint. Thus, the opposing FAD shifts in superior versus inferior regions provide a calibration-independent redox signature that reinforces the superior-to-inferior energetic gradient revealed by both cyto-iATP and mito-iATP measurements. We next asked whether this functional and redox hierarchy is underpinned by a corresponding gradient in mitochondrial capacity.

### Regional differences in mitochondrial abundance define distinct SA node myocyte phenotypes

The opposing FAD redox shifts in superior versus inferior regions provide a calibration-independent signature that reinforces the energetic gradient revealed by cyto-iATP and mito-iATP. A parsimonious explanation for these coupled functional differences is a structural one: the superior SA node may be endowed with greater mitochondrial capacity, enabling higher beat-locked ATP throughput and a distinct redox operating point. Thus, we next asked whether this functional and redox hierarchy is underpinned by a corresponding gradient in mitochondrial capacity, testing the hypothesis that mitochondrial abundance scales with the superior–inferior energetic hierarchy. To this end, we quantified mitochondrial content, first in intact-node whole mounts and then in anatomically identified single myocytes (Figure 7).

**Figure 7.**
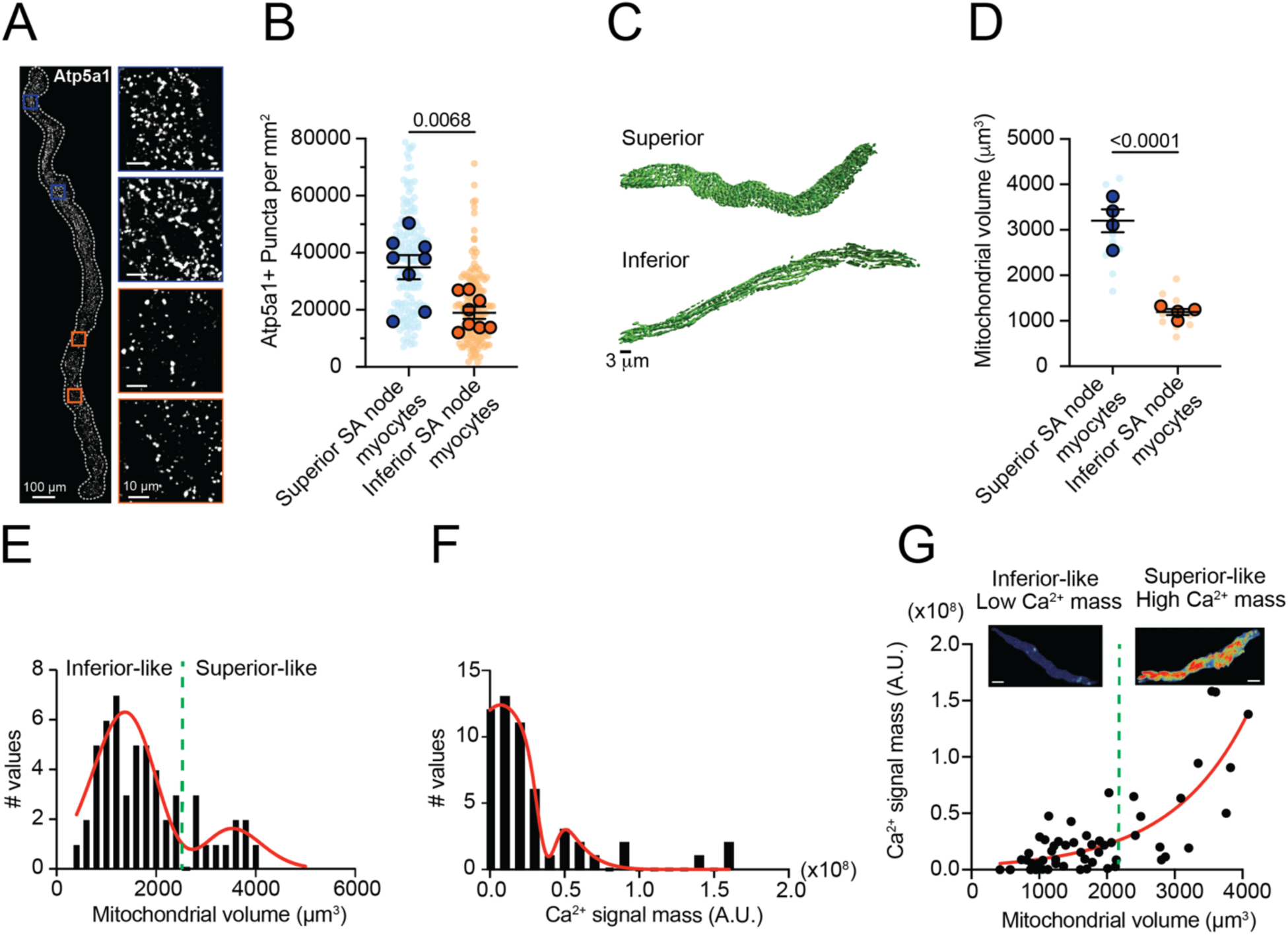
Regional heterogeneity in mitochondrial volume and Atp5a1 expression in SA node myocytes. (**A**) Representative RNAscope in situ hybridization image of a whole-mount SA node labeled for Atp5a1 mRNA (white). Insets show higher-magnification views of Atp5a1 puncta in superior (blue outlines) and inferior (orange outlines) regions. Scale bars: 100 µm (whole mount), 10 µm (insets). (**B**) Quantification of Atp5a1 puncta density in superior and inferior SA node myocytes. (**C**) Representative three-dimensional surface renderings of the mitochondrial reticulum (MitoTracker Green) in myocytes isolated from superior (top) and inferior (bottom) regions. (**D**) Quantification of total mitochondrial volume per myocyte (N = 4 mice per group). (**E**, **F**) Frequency distributions of mitochondrial volume (**E**) and integrated Ca^2+^ signal mass (**F**) pooled across all analyzed myocytes. Red curves indicate Gaussian fits, resolving two populations. The green dashed line (≈ 2.2 × 10^3^ µm^3^) denotes the intersection used to classify low- and high-volume/mass phenotypes. (**G**) Relationship between mitochondrial volume and Ca^2+^ signal mass for individual myocytes. The red curve represents an exponential fit (R^2^ = 0.73). Insets show maximum-intensity projections of representative Ca^2+^ transients from low- and high-volume groups. Scale bars, 5 µm. *P*-values are shown above comparisons. Large circles denote per-animal means; small circles indicate individual biological replicates.

To assess whether regional energetic differences have a structural basis, we used *in situ* hybridization (RNAscope) to measure Atp5a1 mRNA expression in whole-mount SA node preparations. Atp5a1 encodes the catalytic α-subunit of F1Fo-ATP synthase (Walker, 2013; Kühlbrandt, 2019), the enzyme responsible for mitochondrial ATP production. The superior region exhibited visibly denser Atp5a1 labeling than the inferior region (Figure 7A). Quantification confirmed a significantly higher density of Atp5a1-positive puncta in the superior domain (*P* = 0.0068; Figure 7B). This mRNA gradient suggests that superior SA node cells are transcriptionally programmed to produce more ATP synthase, potentially supporting greater oxidative phosphorylation capacity. To complement this molecular analysis, we performed volumetric 3D reconstructions of region-identified, isolated SA node myocytes labeled with MitoTracker Green to assess total mitochondrial mass. Superior cells possessed substantially larger mitochondrial networks than inferior cells (Figure 7C), with a >2-fold increase in total mitochondrial volume (3014 ± 245 µm^3^ vs 1161 ± 121 µm^3^; *P* < 0.0001; Figure 7C**, D**). Thus, superior SA node myocytes benefit from both increased mitochondrial mass and elevated Atp5a1 expression, suggesting convergent structural and biochemical mechanisms underlying their high-gain ATP phenotype.

To determine whether mitochondrial size constrains the Ca^2+^ burden of individual pacemaker cells, we analyzed the full pool of SA node myocytes instead of pre-sorting them by anatomical origin. Pooling was justified for three reasons: (i) 3D reconstructions revealed a clearly bimodal distribution of mitochondrial volumes—small-volume cells came almost exclusively from the inferior node and large-volume cells from the superior node—so treating the tissue as two rigid classes would have narrowed the quantitative range available for correlation analysis and reduced power. (ii) Ca^2+^ transients fire at ∼4 Hz in superior myocytes versus ∼1 Hz in inferior cells (Grainger et al. (2021) because transient amplitudes are comparable, frequency alone accounts for their greater total Ca^2+^ flux. In our dataset, every Ca^2+^ record was normalized to its own firing rate, removing this regional bias and allowing us to test whether bigger mitochondrial networks support larger beat-normalized Ca^2+^ loads across the entire spectrum of phenotypes. (iii) The limited yield of viable, dye-loaded cells obtained from finely dissected sub-regions—and the need to collect both 3D mitochondrial stacks and high-speed Ca^2+^ movies—would have left separate regional datasets under-powered. Pooling, therefore, maximizes sample size and provides an unbiased, statistically robust test of structure–function coupling.

3D mitochondrial stacks, obtained by SRRF imaging of the mitochondrial reticulum, and movies of cytosolic Ca^2+^ signals were generated from acutely dissociated SA node myocytes loaded with MitoTracker and Fluo-4 AM. Mitochondrial-volume histograms displayed two well-separated peaks; a double-Gaussian fit yielded a threshold of ∼2,200 µm^3^ that segregated “inferior-like” (low-volume) from “superior-like” (high-volume) cells (Figure 7E). An analogous bimodal pattern was observed for Ca^2+^ signal mass (Figure 7F). Across the pooled population, mitochondrial volume was a strong predictor of Ca^2+^-handling load: larger mitochondria supported proportionally greater Ca^2+^ signal mass, with an exponential fit explaining 73% of the variance (Figure 7G). Ca^2+^ signal mass scaled with mitochondrial volume, such that cells with lower mitochondrial content failed to reach the Ca^2+^ output observed in high-volume cells, suggesting that organelle size constrains the upper limit of signaling capacity.

Together with the higher beat-locked cyto- and mito-iATP throughput in the superior node (see Figures 1–3, above) and the faster intrinsic firing previously reported in superior versus inferior SA node myocytes (Grainger et al., 2021), these data indicate that superior pacemaker cells combine an expanded mitochondrial network with stronger Ca^2+^ signaling and higher energetic throughput, whereas inferior cells couple reduced mitochondrial content to lower Ca^2+^-linked demand and lower ATP turnover. Thus, regional differences in mitochondrial volume represent a structural determinant that helps define distinct metabolic and electrophysiological phenotypes within the SA node (Santana and Earley, In press).

### Beat-locked Ca^2+^ transients trigger cytosolic ATP increases in SA node myocytes

Having shown that mitochondrial volume scales with the Ca^2+^ burden that a cell must service, we next asked whether the timing of Ca^2+^ release likewise governs the beat-to-beat rise of ATP in the cytosol, turning structural capacity into real-time metabolic output. To do this, we loaded dissociated SA node myocytes expressing cyto-iATP with the red-shifted Ca^2+^ indicator, Rhod-3, and simultaneously tracked cytosolic ATP and Ca^2+^ using a confocal microscope in line-scan mode (Figure 8A).

**Figure 8.**
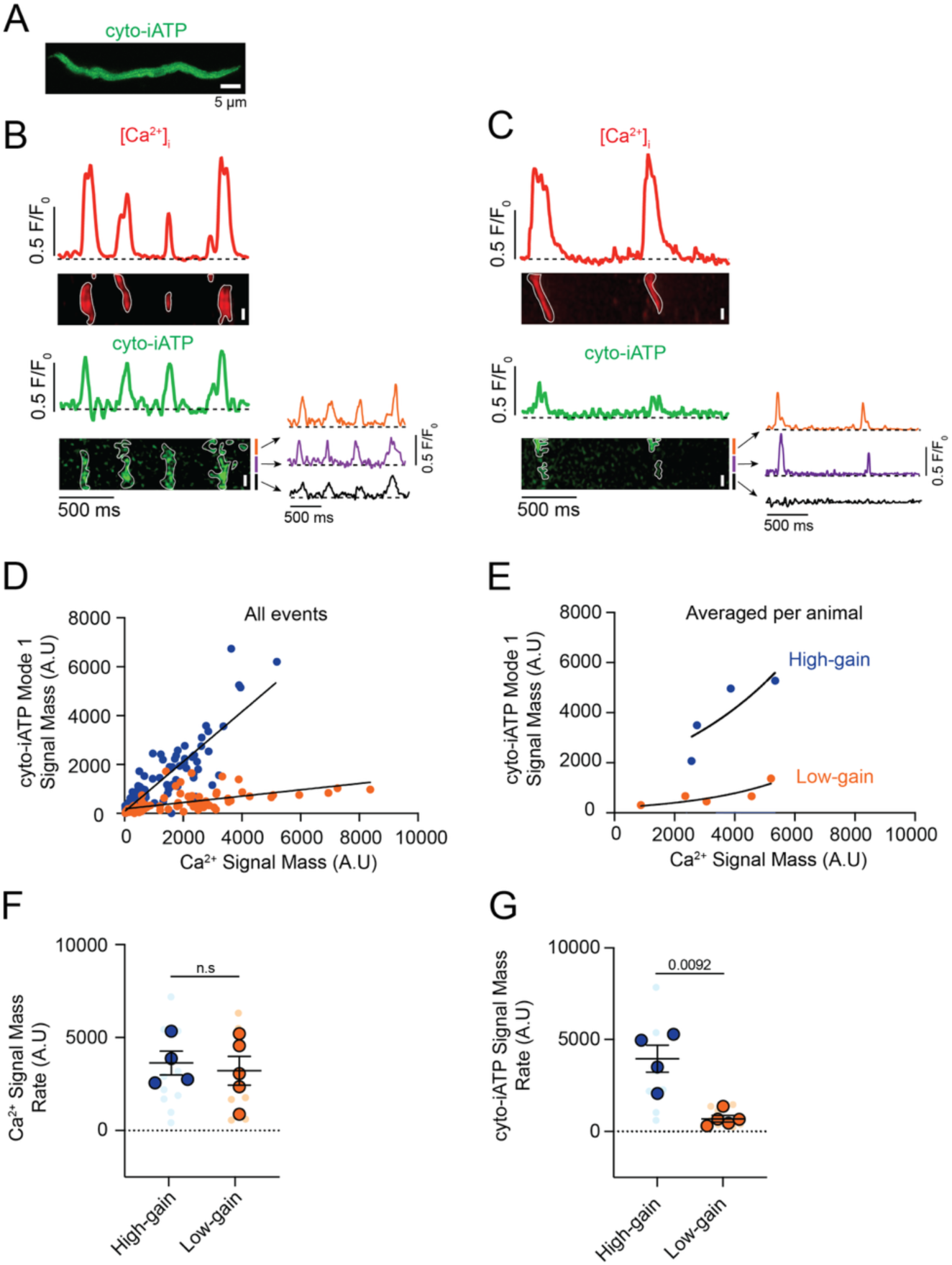
Beat-to-beat ATP synthesis is coupled to intracellular Ca^2+^ release through distinct high- and low-gain metabolic transfer functions in SA node myocytes. (**A**) Representative confocal image of an isolated SA node myocyte expressing the cytosolic ATP sensor (cyto-iATP). Scale bar, 5 µm. (**B**, **C**) Simultaneous confocal line-scan recordings of intracellular Ca^2+^ (red, top) and cyto-iATP (green, bottom). (**B**) Example of a myocyte classified as high-gain, showing temporally aligned Ca^2+^ transients and cyto-iATP signals. Traces extracted from local regions of interest (colored traces) illustrate spatially resolved Ca^2+^ and ATP dynamics. (**C**) Example of a myocyte classified as low-gain, showing Ca^2+^ transients with minimal corresponding cyto-iATP signals. (**D**) Event-level relationship between Ca^2+^ signal mass (input) and cyto-iATP signal mass (output). Data are separated into two populations: high-gain events (blue circles) and low-gain events (orange circles). Solid lines indicate linear fits for each population. (**E**) Cell-averaged relationship between Ca^2+^ and cyto-iATP signal mass, preserving bimodal separation between high-gain (blue) and low-gain (orange) groups. (**F**, **G**) Summary quantification of Ca^2+^ signal mass rate (**F**) and cyto-iATP signal mass rate (**G**) for high- and low-gain populations. *P*-values are shown above comparisons. Large circles denote per-animal means; small circles indicate individual biological replicates.

Simultaneous line-scan imaging of Ca^2+^ release and cyto-iATP fluorescence in two representative pacemaker cells with high (Figure 8B) and low (Figure 8C) frequency of Ca^2+^-release events showed that, in both cases, every Ca^2+^-release event is paired with a rise in cytosolic ATP, confirming beat-locked metabolic coupling. The spatial extent of the ATP rise was found to differ markedly: in the high-frequency cell (Figure 8B), the increase was spread across a much broader swath of the cytoplasm, whereas in the low-frequency cell (Figure 8C), the increase was restricted to a narrow segment. Quantitative differences in these patterns, exemplified by traces from three regions of interest shown beneath each cyto-iATP image, highlight localized versus widespread ATP transients.

Given the variations in spatial distributions, we quantified the signal mass of Ca^2+^ signals and their associated cyto-iATP events. As shown in Figure 8D**, E** pooled events from all cells imaged segregated into two groups, both of which could be fit to linear functions distinguished by the different slopes of their Ca^2+^-ATP relationships: 1.02 Δcyto-iATP/ΔCa^2+^ for high-gain events and 0.12 Δcyto-iATP/ΔCa^2+^ for low-gain events (*P* < 0.0001). Notably, plotting averaged cyto-iATP and Ca^2+^ signal mass per cell segregated the data into two clear gain regimes: a low-gain group with minimal ATP amplification and a high-gain group with steep, exponential Ca^2+^-to-ATP coupling (Figure 8E).

Despite the similarity of Ca^2+^ signal-mass rate (Figure 8F) across phenotypes, cyto-iATP signal-mass rate was significantly larger in high-gain cells (*P* = 0.0092), underscoring the higher energetic output in these cells over time (Figure 8G). This dichotomy mirrors the intact node, where superior myocytes exhibit greater signal-mass rates than inferior myocytes; it is further reinforced by higher event frequencies in high-gain cells (2.56 ± 0.16 Hz) versus low-gain cells (1.50 ± 0.18 Hz; *P* = 0.003) that closely match regional values in tissue (Figures 1**, 2**).

A kinetic analysis of Ca^2+^ transients showed slower dynamics in high-gain cells, with a longer decay time (t_1/2_ = 92.48 ± 20.77 ms; *P* = 0.0419) and prolonged duration (434.70 ± 50.83 ms; *P* = 0.0125) compared with low-gain cells (t_1/2_ = 41.11 ± 8.70 ms; 247.40 ± 33.71 ms) (**Figure S4A–E**). In contrast, cyto-iATP transients exhibited comparable rise times in high-gain (99.81 ± 10.56 ms) and low-gain (90.88 ± 11.30 ms) cells, as well as similar decay times (t_1/2_ ∼47–66 ms) and durations (∼247.4–286.1 ms) (**Figure S4F–H**). Thus, cytosolic ATP output in pacemaker myocytes is not simply proportional to Ca^2+^ load but instead falls into discrete high- and low-gain modes, echoing the superior–inferior metabolic hierarchy of the intact SA node.

### High- and low-gain mitochondrial ATP production in SA node myocytes

We found that Mode 1 events—rapid post-beat rises in mito-iATP—are tightly beat-locked, with Ca^2+^ transients preceding ATP onset by <50 ms, consistent with rapid SR-to-mitochondria Ca^2+^ transfer gating oxidative phosphorylation. The magnitude of ATP elevation per unit Ca^2+^ load varied sharply between cells. Representative high-gain cells (Figure 9A) produced broad, high-amplitude ATP increases spanning most of the scan region, while low-gain cells (Figure 9B) generated smaller, spatially restricted rises, despite comparable Ca^2+^ signals.

**Figure 9.**
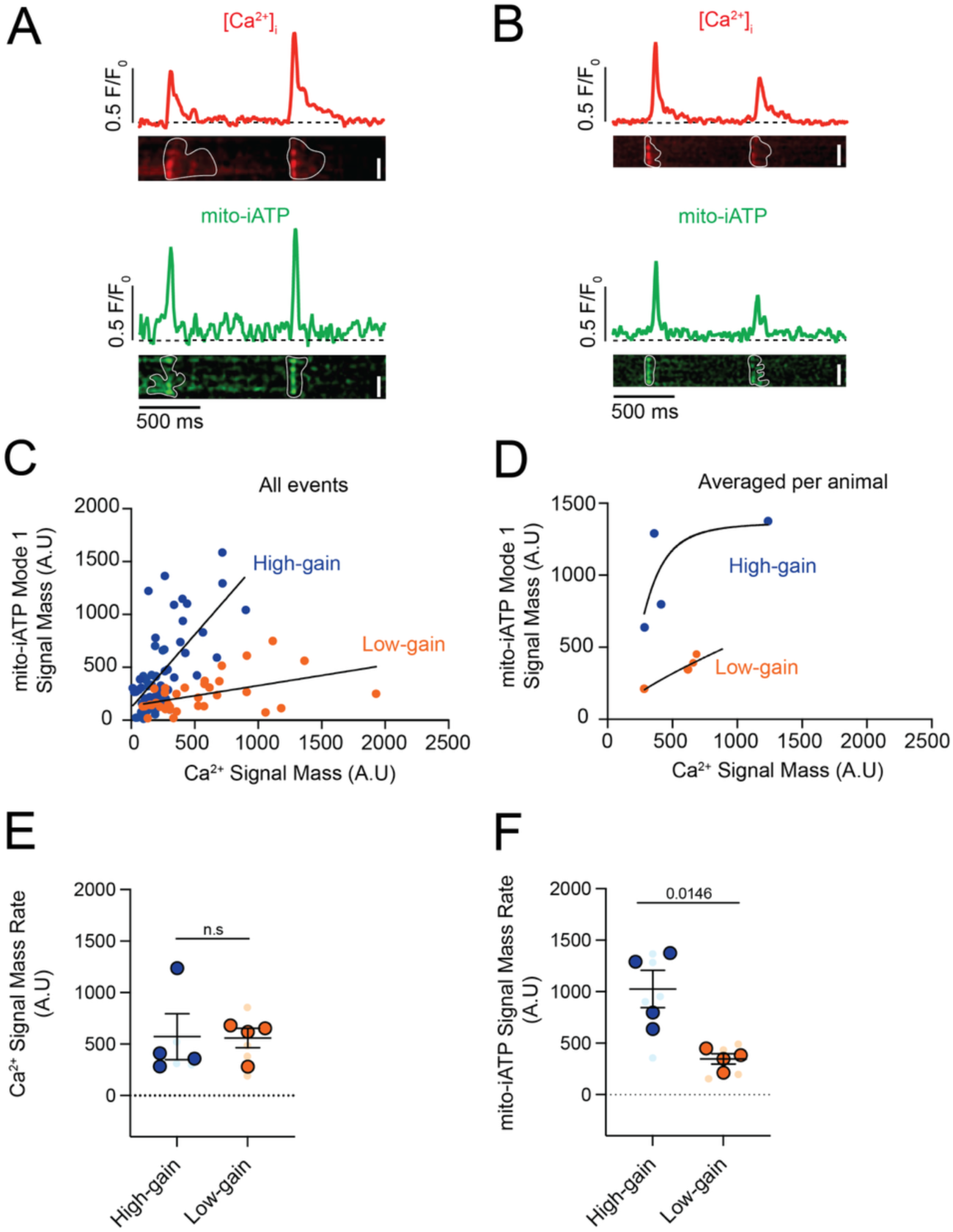
Beat-to-beat mitochondrial ATP synthesis exhibits bimodal coupling to intracellular Ca^2+^ release in SA node myocytes. (**A**, **B**) Representative confocal line-scan recordings showing simultaneous intracellular Ca^2+^ transients (red, top) and mitochondrial ATP signals (mito-iATP; green, bottom). (**A**) Example of a myocyte classified as high-gain, displaying temporally aligned Ca^2+^ transients and mito-iATP signals. (**B**) Example of a myocyte classified as low-gain, in which Ca^2+^ transients are accompanied by minimal mito-iATP responses. (**C**) Event-level relationship between Ca^2+^ signal mass (input) and mito-iATP signal mass (output). Data is segregated into two populations: high-gain events (blue circles) and low-gain events (orange circles), with solid lines indicating linear fits. (**D**) Cell-averaged relationship between Ca^2+^ and mito-iATP signal mass, fitted with a Hill function for each group (high-gain, blue; low-gain, orange). (**E**, **F**) Summary quantification of Ca^2+^ signal mass rate (**E**) and mito-iATP signal mass rate (**F**) for high- and low-gain populations. *P*-values are shown above comparisons. Data are presented as means ± SEM (N = 4 mice per group). Large circles denote per-animal means; small circles indicate individual biological replicates.

Pooling the signal mass of all associated Ca^2+^ and mito-iATP events across cells yielded an apparently piecewise-linear Ca^2+^–ATP relation with two regimes: low slope (0.19 Δmito-iATP/ΔCa^2+^) and high slope (1.38 Δmito-iATP/ΔCa^2+^) (Figure 9C); but because events are nested within cells, this marginal pattern reflects aggregation. Accordingly, per-cell fits showed Hill models with greater apparent cooperativity in high-gain cells (Hill coefficient, 2.88) than low-gain cells (Hill coefficient, 0.78) and distinct plateaus (Figure 9D). The sigmoidal behavior is expected from thresholded, cooperative mitochondrial Ca^2+^ control, where MICU1/2 gatekeeping of the MCU sets the Ca^2+^ threshold and gain (Mallilankaraman et al., 2012; Csordas et al., 2013; Kamer et al., 2017). Plateau differences imply capacity differences: the lower maximum output (ceiling) in low-gain cells is consistent with reduced mitochondrial mass and/or diminished per-mitochondrion respiratory capacity. It is important to note, however, that because iATPSnFR family sensors have finite dynamic ranges and can saturate at high ATP (Lobas et al., 2019; Marvin et al., 2024; Rhana et al., 2024), the upper plateau in high-gain cells could partly reflect sensor saturation rather than biology, so ceiling differences are interpreted cautiously.

We extended the analysis by calculating Ca^2+^ and mito-iATP signal-mass rates (total signal mass per trace divided by recording time), which index Ca^2+^ load and ATP production over time rather than per-event yield. Ca^2+^ signal-mass rate did not differ between groups (Figure 9E), whereas mito-iATP signal-mass rate was higher in high-gain cells (*P* = 0.0146) (Figure 9F). Per-cell averages corroborated the two phenotypes, with high-gain cells firing faster (2.49 ± 0.59 Hz vs 0.92 ± 0.12 Hz; *P* = 0.0164). For Ca^2+^ transients, rise times did not differ significantly (106.40 ± 17.28 ms vs 70.39 ± 7.23 ms), but high-gain cells exhibited slower decay (t_1/2_ = 85.31 ± 15.65 ms vs 39.63 ± 12.17 ms) and longer duration (421.60 ± 65.36 ms vs 251.90 ± 46.28 ms) (**Figure S5A, C–E**). Mode 1 Mito-iATP event kinetics were similar between groups (rise time 94.40 ± 17.39 ms vs 67.11 ± 14.70 ms; decay t_1/2_, ∼59–64 ms; duration ∼299.80–301.0 ms) (**Figure S5B, F–H**).

Thus, Mode 1 mitochondrial ATP production is stratified into low- and high-gain phenotypes with shallower versus steeper Ca^2+^–ATP gain; high-gain cells rapidly mobilize ATP once a Ca^2+^ gate is crossed and operate nearer a higher ceiling, whereas low-gain cells exhibit a lower ceiling and approach saturation only at higher Ca^2+^ loads. This functional dichotomy echoes the superior–inferior metabolic hierarchy of the intact SA node, revealing that intrinsic cooperativity shapes not only the magnitude but also the dynamic range of pacemaker cell bioenergetics.

To provide a preparation-matched control for optical artifacts in isolated myocytes—and to complement the intact-node measurements obtained in the presence of blebbistatin—we co-expressed mito-iATP with an ATP-insensitive HaloTag reporter labeled with the HaloTag Ligand, Janelia Fluor JFX554, and imaged both channels simultaneously. Mito-iATP fluorescence localized to a punctate mitochondrial network, whereas JFX554-HaloTag labeling was spatially stable across the same cell (**Figure S6A**). In representative Mode 1 recordings, beat-locked mito-iATP transients were robust, while the HaloTag channel remained flat; the Mode 1 waveform was preserved by ratiometric normalization (mito-iATP/HaloTag) (**Figure S6B**). Across cells, the ratiometric Mode 1 peak amplitude tracked the raw mito-iATP peak amplitude with strong linearity (R^2^ = 0.9402; **Figure S4C**), indicating that the oscillations are not explained by motion, drift, or field-stimulation–associated fluorescence fluctuations. The same held for Mode 2 responses: mito-iATP dips occurred without corresponding changes in the HaloTag channel, and the mito-iATP/HaloTag ratio preserved Mode 2 events (**Figure S6D**), with ratiometric amplitudes closely matching the raw mito-iATP dip amplitudes (R^2^ = 0.9920; **Figure S6E**). Collectively, these panel-resolved controls strengthen the conclusion that beat-locked mito-iATP waveforms reflect genuine ATP dynamics rather than movement- or optics-driven artifacts.

### High- and low-load mitochondrial ATP consumption in SA node myocytes

As was the case in intact SA nodes, we detected Mode 2 events—transient mito-iATP dips following each Ca^2+^ transient, indicating rapid, beat-synchronous ATP depletion in the matrix mitochondria—in isolated myocytes (Figure 10). Plotting mito-iATP dip signal mass versus Ca^2+^ signal mass (**Figure 10A, B**) revealed two regimes separated by an empirical Ca^2+^ threshold of ∼2,000 AU: below this threshold, events formed a dense, shallow-dip cluster (slope = -0.07 Δmito-iATP/ΔCa^2+^), whereas above it, dips deepened with increasing Ca^2+^ load (slope = -0.26 Δmito-iATP/ΔCa^2+^), as summarized in **Figure 10C, D**. High-load cells drew down ∼2-fold more mitochondrial ATP per beat than low-load cells (-905.92 ± 162.30 vs. -407.40 ± 82.41 AU) and fired faster (2.11 ± 0.20 vs. 1.08 ± 0.23 Hz). For Ca^2+^ transients, kinetics did not differ between groups (rise, 80.88–90.01 ms; t_1/2_, 32.88–46.69 ms; duration, 230.90–267.60 ms; **Figure S7A–E**). Similarly, Mode 2 mito-iATP kinetics did not differ between groups (rise, 91.33–107.30 ms; t_1/2_, 39.70–51.21 ms; duration, 232.90–304.60 ms; **Figure S7F–H**).

**Figure 10.**
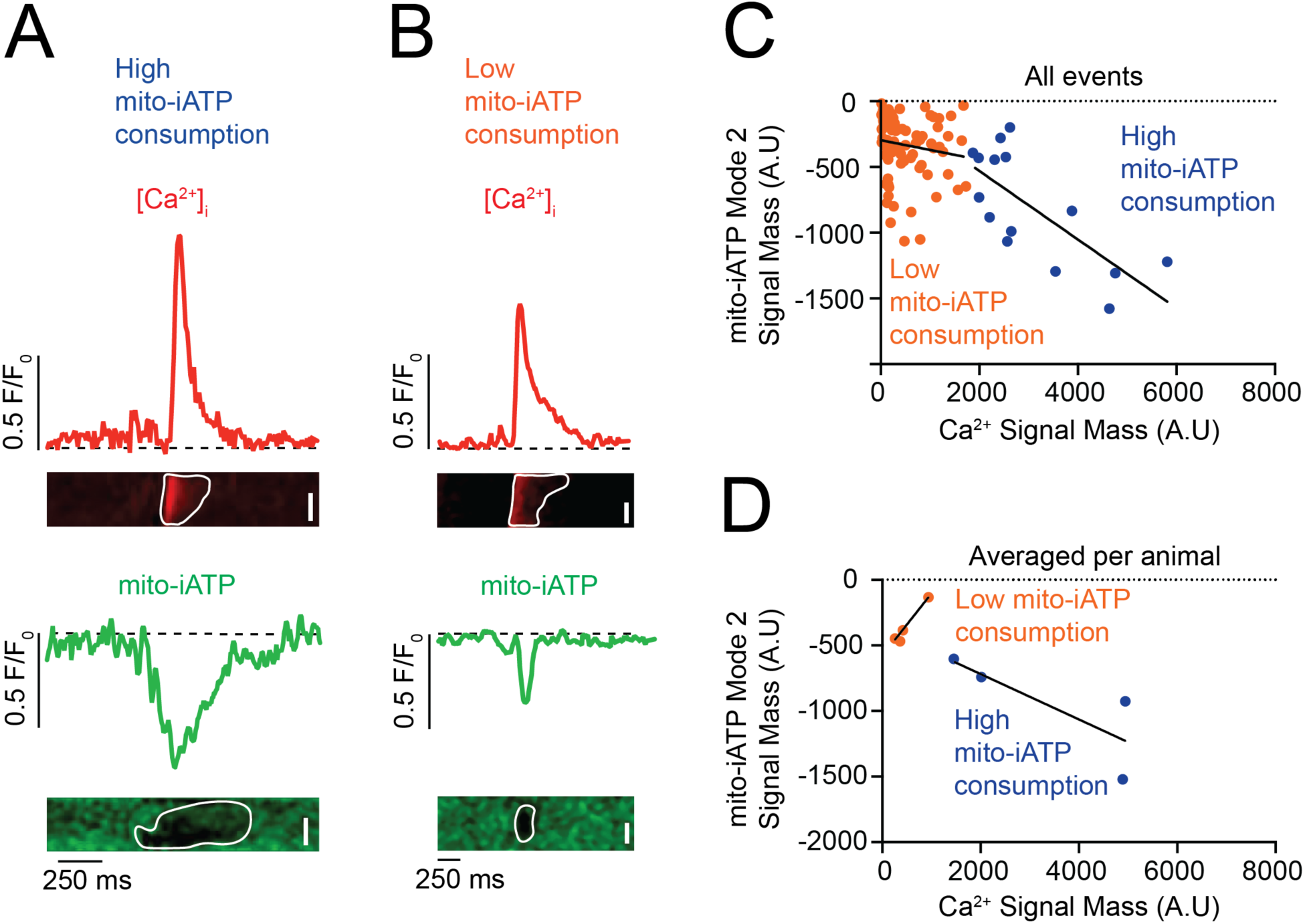
Beat-to-beat mitochondrial ATP consumption exhibits bimodal coupling to intracellular Ca^2+^ release in SA node myocytes. (**A**, **B**) Representative confocal line-scan recordings showing simultaneous intracellular Ca^2+^ transients (red, top) and Mode 2 mitochondrial ATP signals (mito-iATP; green, bottom). (**A**) Example of a myocyte classified as high-load, in which Ca^2+^ release is associated with large-amplitude negative mito-iATP deflections. (**B**) Example of a myocyte classified as low-load, showing Ca^2+^ transients with smaller accompanying mito-iATP deflections. (**C**) Event-level relationship between Ca^2+^ signal mass (input) and mitochondrial ATP consumption signal mass (output). Data is segregated into two populations: high-load events (blue circles) and low-load events (orange circles). Solid lines indicate linear fits for each population. (**D**) Cell-averaged relationship between Ca^2+^ signal mass and mitochondrial ATP consumption for high-load (blue) and low-load (orange) groups. Scale bars, 5 µm. Data are presented as mean fits (N = 4 mice per group). In (**C**), circles represent individual biological events; in (**D**), circles denote per-animal means.

Thus, mitochondrial ATP consumption is stratified into low- and high-load phenotypes: cells with mito-iATP dips above ∼2,000 AU consume disproportionately more ATP per beat, whereas cells below this threshold consume less despite similar Ca^2+^ inputs, indicating discrete utilization modes that likely contribute to metabolic specialization within the node.

## Discussion

In this study, we demonstrate that ATP production in SA node myocytes is synchronized to each heartbeat rather than being produced continuously and is confined to discrete metabolic microdomains. By imaging mitochondrial ATP alongside Ca^2+^ transients, we observed recurring “ATP gain” peaks and “ATP dip” troughs whose magnitudes and timing matched the electrical cycle, revealing a just-in-time energetic budget tailored to beat-by-beat ionic demand.

### ATP microdomains and regional hierarchy

Regional analyses uncovered pronounced heterogeneity along the node. Superior SA node myocytes, which possess greater mitochondrial volume and reside in regions of higher capillary density, displayed larger ATP gains and higher diastolic [ATP]_i_ than inferior cells, in line with their faster basal firing rates. Pharmacological experiments illuminated SR-to-mitochondria Ca^2+^ transfer as the proximate trigger for oxidative phosphorylation: thapsigargin abolished ATP oscillations by preventing SR Ca^2+^ refilling, whereas blocking depolarizing HCN channels with ivabradine simply slowed their frequency without affecting their amplitude, demonstrating that the SR Ca^2+^ cycling governs metabolic activation upstream of the I_f_-dependent membrane depolarization.

These structure–function relationships extend earlier anatomical work showing a superior-to-inferior vascular gradient in the mouse node (Grainger et al., 2021). Manning et al. (2024) linked chronic under-perfusion- to hypoxia-induced factor (HIF)–-1α activation, SERCA down-regulation, and reduced PGC-1α–dependent mitochondrial biogenesis. Our results place this hypoxia–HIF cascade within the nodal context, supporting the conclusion that microvascular “wealth” establishes the mitochondrial “capital” available for rapid ATP payouts.

The dominant pacemaking locus within the SA node is also dynamic, migrating along the superior–inferior axis in response to changes in autonomic tone (Yamamoto et al., 1998; Boyett et al., 2000; Brennan et al., 2020). Optical mapping of *ex vivo* rat and human preparations demonstrates the coexistence of two discrete initiation sites—a richly vascularized, mitochondria-dense superior focus and a comparatively hypoperfused inferior focus—that alternate leadership according to sympathetic versus parasympathetic drive (Brennan et al., 2020). Our findings place firm energetic limits on this plasticity: when initiation shifts toward the inferior node, the lower capillary density, reduced mitochondrial reserve, and smaller beat-to-beat ATP gains of this region constrain the maximal rate it can sustain. Thus, although the pacemaker center can relocate, the performance of the newly recruited site remains bounded by the surrounding microvascular and metabolic landscape that we have mapped.

### Metabolic coupling or MCU-ANT axis

Our analysis revealed two coupling modes in pacemaker myocytes. Mode 1 is characterized by beat-locked mitochondrial ATP “gains” and exhibits two phenotypes: a high-gain subtype in which modest Ca^2+^ transients elicit steep ATP increases, and a low-gain subtype with a shallower Ca^2+^ -to-ATP coupling slope. Mode 2 shows mitochondrial ATP “dips” that scale linearly and inversely with Ca^2+^ load, consistent with consumption-dominated control. Together, these phenotypes underscore heterogeneity in SR–mitochondrial communication, likely reflecting differences in Ca^2+^-transfer efficacy, mitochondrial reserve, and local substrate availability.

The RU360 and BKA experiments provide direct mechanistic evidence that these ATP modes arise from a bidirectional molecular apparatus rather than passive buffering: both mitochondrial and cytosolic ATP transients are abolished by blocking either MCU or ANT, while Ca^2+^ cycling persists. The elimination of Mode 1 ATP gains by RU360 shows that the gain phase is actively driven by Ca^2+^ entry through the MCU, whereas ANT block demonstrates that beat-locked ATP pulses vanish if ADP cannot enter—and ATP cannot exit—the matrix. In this framework, the Ca^2+^ clock provides the trigger, MCU and ANT implement the high-gain metabolic response, and the myocyte falls back onto basal glycolytic and phosphocreatine reserves when this axis is disrupted, transitioning from a “superior-like”, high-bandwidth state to an “inferior-like”, low-bandwidth state in which Ca^2+^ cycling continues but is energetically weakened.

While our experiments establish the existence of high- and low-gain ATP responses, they do not resolve the underlying molecular determinants. We therefore propose two testable hypotheses for future work: 1) a structure-centric model based on differential SR–mitochondria tethering, and 2) a biochemical framework centered on variable oxidative phosphorylation capacity. In the first case, superior pacemaker cells may express more Mfn2 (or other tethers), narrowing the SR-mitochondrial cleft and amplifying beat-triggered Ca^2+^ transfer. In this scenario, inferior cells, with sparser coupling, would receive smaller mitochondrial Ca^2+^ pulses and thus operate at lower metabolic gain. In the second case, high-gain cells would harbor a denser complex-V proteome (e.g., greater ATP5A content), allowing faster ATP synthesis per unit Ca^2+^. In this scenario, low-gain cells, with reduced ATP5A abundance, would be constrained to shallower slopes. These structural (coupling) and biochemical (flux capacity) gradients together offer a plausible framework for the dual-gain behavior we observe and define a roadmap for future mechanistic dissection. Together, these structural (coupling) and biochemical (flux capacity) gradients offer a plausible framework for the dual-gain behavior we observe and define a roadmap for future mechanistic dissection.

Our RNAscope analysis provides direct molecular validation of the biochemical hypothesis, revealing significantly higher Atp5a1 mRNA expression in superior versus inferior SA node regions. ATP5A1 encodes the catalytic α-subunit of F1Fo-ATP synthase (Walker, 2013; Kühlbrandt, 2019), and elevated transcript levels likely translate to increased enzyme abundance and enhanced ATP synthetic capacity per mitochondrion. This transcriptional gradient is mechanistically significant for several reasons. First, the Atp5a1 difference may exceed the 2.6-fold mitochondrial volume gradient we observe between superior and inferior cells, indicating convergent structural and biochemical advantages: superior myocytes benefit from both increased mitochondrial mass and higher ATP synthase content per organelle, compounding their energetic capacity. Second, the mRNA gradient reveals transcriptional programming rather than acute metabolic adjustment, implicating upstream regulators such as PGC-1α and estrogen-related receptor α (ERRα)—which are themselves oxygen-sensitive and vascular supply-dependent—as determinants of regional pacemaker identity (Rowe et al., 2010; Scarpulla, 2011). This connects our molecular findings to the vascular heterogeneity reported by (Grainger et al., 2021) and suggests that superior cells maintain higher Atp5a1 expression because they receive better oxygen delivery, establishing a feed-forward loop in which vascular supply drives mitochondrial biogenesis and ATP synthase content. Our findings complement recent spatial transcriptomics studies identifying oxidative phosphorylation gene heterogeneity in cardiac conduction system cells (Liang et al., 2021; Kanemaru et al., 2023) by demonstrating that metabolic programming creates functional gradients within SA node regions. Together, these findings establish that the high-gain ATP phenotype in superior cells has a transcriptional basis: enhanced vascular oxygen delivery (Grainger et al., 2021) drives PGC-1α/ERRα-dependent upregulation of Atp5a1 (Rowe et al., 2010; Scarpulla, 2011), amplifying ATP synthetic capacity through both increased mitochondrial mass and elevated ATP synthase content per organelle.

### SA node heterogeneity in the context of entrainment and stochastic resonance

It is important to place our findings in the context of two classical synchronization mechanisms that operate in oscillatory biological networks: entrainment and stochastic resonance. Entrainment refers to the mutual electrotonic coupling of self-oscillating myocytes, whereby the cell or region with the shortest intrinsic cycle length imposes its rhythm on its neighbors, achieving network-wide phase alignment (Jalife, 1984; Anumonwo et al., 1991). This process is most efficient when all cells possess comparable metabolic reserves, as would be expected in a node endowed with uniformly high vascular density that affords each myocyte abundant mitochondrial ATP production. Stochastic resonance, in contrast, is a nonlinear phenomenon in which intrinsic noise enhances the detection or propagation of weak periodic signals (Wiesenfeld and Moss, 1995; Gammaitoni et al., 1998; Hanggi, 2002). In the SA node, such noise can originate from metabolically constrained, intermittently firing cells situated in regions of sparse perfusion (Grainger et al., 2021; Guarina et al., 2022). When microvascular rarefaction creates metabolic heterogeneity, the resulting juxtaposition of high-gain and low-gain myocytes can harness stochastic resonance, preserving overall heart-rate frequency and periodicity, even as localized pockets of low excitability emerge.

A growing literature demonstrates pronounced electrophysiological heterogeneity along the SA node (Boyett et al., 2000). High-resolution 3D imaging showed discontinuously and asynchronously propagating Ca^2+^ transients in the intact mouse SA node, arising from spatially heterogeneous local Ca^2+^ release events within the HCN4^+^/Cx43^-^ meshwork (Kim et al., 2018; Monfredi et al., 2018; Bychkov et al., 2020). Complementary work revealed parallel heterogeneity in vascular density, membrane excitability, and Ca^2+^ signaling along the longitudinal axis of the node (Grainger et al., 2021). Such variability motivated the hypothesis that intrinsic noise sharpens pacemaker timing via stochastic resonance (Clancy and Santana, 2020; Grainger et al., 2021; Guarina et al., 2022). A recent multiscale study confirmed that such noise fine-tunes SA node firing and preserves rhythmicity under strong parasympathetic drive (Okamura et al., 2024). Collectively, these data establish microscopic heterogeneity—spanning structure, metabolism, and electrical behavior—as an intrinsic feature of native SA node tissue that critically modulates its function.

Our current findings provide a metabolic explanation for this variability: inferior SA node myocytes, constrained by sparse vascular supply and limited mitochondrial reserve, behave as low-frequency, intermittently firing “noise generators.” Observations of beat-locked mitochondrial ATP dips in Mode 2 cells confirm that, because of their half-recovery time of ∼125 ms, these dip-dominant cells cannot entrain to the rapid rates sustained by gain-dominant neighbors at 8–10 Hz. Instead, the pronounced variability in ATP-withdrawal and -recovery kinetics creates the metabolic noise required for stochastic resonance, such that subthreshold fluctuations in membrane potential driven by deep ATP dips enhance the sensitivity of electrotonically coupled networks to depolarizing inputs from well-perfused, high-gain cells. Importantly, the failure of dip-dominant cells to follow 8–10 Hz pacing, revealed by beat-locked mito-iATP dips, and their subsequent noise-generating behavior, provides the first metabolic assay capable of distinguishing entrainment from stochastic resonance. Our model predicts maximal network coherence when these low-gain, noise-generator cells are coupled to high-gain, well-perfused superior myocytes; this aligns with our experimental finding that collapsing mitochondrial reserve with FCCP or thapsigargin abolishes resonance, underscoring its metabolic origin.

While our experiments were performed in the murine SA node, the concepts we propose have broad applicability, as the bioenergetic machinery we interrogate—mitochondrial Ca^2+^ uptake, tricarboxylic acid cycle (TCA) cycle activation, and oxidative phosphorylation—are widely conserved. Thus, the insights gained in murine models likely translate broadly, including to human cardiac physiology. Consistent with this premise, Tagirova Sirenko et al. (2021) showed that the fundamental pacemaking architecture of mouse and human SA node cells is essentially the same. We therefore anticipate that the electro-metabolic principles revealed here will apply to any excitable tissue or intrinsic pacemaker.

### Capillary-Mitochondria-Ion-Channel axis and clinical implications

Crucially, our SA node findings provide direct support for the Capillary–Mitochondria–Ion-Channel (CMIC) axis recently proposed by our group and colleagues (Santana and Earley, In press), in which excitability emerges from a coupled system in which capillary density sets the local oxygen ceiling, mitochondrial volume defines energetic bandwidth, and ion channels are tuned by metabolic state. Within this framework, [ATP]_i_ measurements link the superior–inferior energetic hierarchy to underlying vascular and mitochondrial architecture such that higher capillary density and shorter myocyte–capillary distances in the superior node align with higher diastolic cytosolic [ATP]_i_ (∼0.8 mM), whereas the more sparsely vascularized inferior node exhibits lower [ATP]_i_ (∼0.4 mM) and more oxidized FAD signals, consistent with a chronically supply-limited state. Microvascular rarefaction can thus be viewed as pushing pacemaker cells “down” the CMIC axis—from a superior-like, well-supplied regime toward an inferior-like, energy-limited regime—so that modest shifts in [ATP]_i_ within the 0.4–0.8 mM window land on the steep portion of the ATP dependence for many metabolic enzymes and ion channels, amplifying the impact of structural vascular changes on beat-to-beat excitability and nodal robustness.

The clinical relevance of this supply-limited phenotype is underscored by a mouse model of early heart failure with preserved ejection fraction (HFpEF) in which selective rarefaction of superior-node capillaries precedes bradycardia and heightened beat-to-beat variability (Manning et al., 2025). We found that rarefaction compromises pacemaker robustness, even when global cardiac output remains normal, indicating that local vascular loss can destabilize rhythm before systemic hemodynamics decline. In the context of this study, however, microvascular rarefaction sculpts pockets of diminished mitochondrial reserve and low excitability, narrowing the roster of regions capable of sustaining high-frequency pacing. When such hypoperfused zones remain interlaced with well-supplied tissue, noise-assisted integration between low-gain and high-gain myocytes can act as a metabolic safety mechanism, injecting timing noise that preserves overall heart-rate frequency and beat-to-beat regularity despite localized deficits. Thus, while the leading pacemaker remains mobile, its performance—and resilience—are ultimately governed by the interplay among vascular architecture, mitochondrial content, and microdomain ATP dynamics. The MCU–ANT axis we uncover here operationalizes this paycheck-to-paycheck logic: each Ca^2+^ transient triggers a burst of oxidative phosphorylation that depends on Ca^2+^ entry into the matrix and on rapid ADP/ATP exchange across the inner membrane.

### Overall Model and Conclusion

In summary, our work extends a unified paycheck-to-paycheck model of cardiac energetics from ventricular myocytes to the pacemaker SA node. In this view, every heartbeat is supported by ATP that is generated, used, and renewed on a cycle-by-cycle basis: ionic and Ca^2+^ clocks set the rate at which ATP is produced; SR-to-mitochondria Ca^2+^ transfer generates ATP via oxidative phosphorylation; and local vascular supply determines the O_2_ supply needed to generate it. Whether a cell initiates the rhythm or follows it, its electrical and mechanical output is underwritten by the same just-in-time metabolic calculus. Variations in capillary density or mitochondrial content, therefore, shape not only contractile power but also the locus and stability of pacemaking. Recognizing that the heart is an organ that functions on the capacity of each beat to generate the ATP it needs for excitation-contraction coupling provides a common framework for interpreting how supply–demand mismatches give rise to arrhythmias, bradycardia, and pump failure across diverse physiological and pathological settings.

## Data availability statement

The raw data files supporting all findings presented in this paper are available from the corresponding author.

## Author contributions

MMC, CM, and LFS conceived and designed the work. MMC, CM, and LFS acquired data for the paper. MMC, CM, PR, DM, GB, DMC, and LFS analyzed data and wrote the paper. All authors have approved the final version of the manuscript and agreed to be accountable for all aspects of the work. All persons designated as authors qualify for authorship, and all those who qualify for authorship are listed.

## Acknowledgements

We thank Dr. Robert Cudmore for helpful discussions. We also thank Michele Persiani and Camryn Sellers-Porter from the McElroy lab for their technical assistance.

---

**Supplemental Figure 1.**
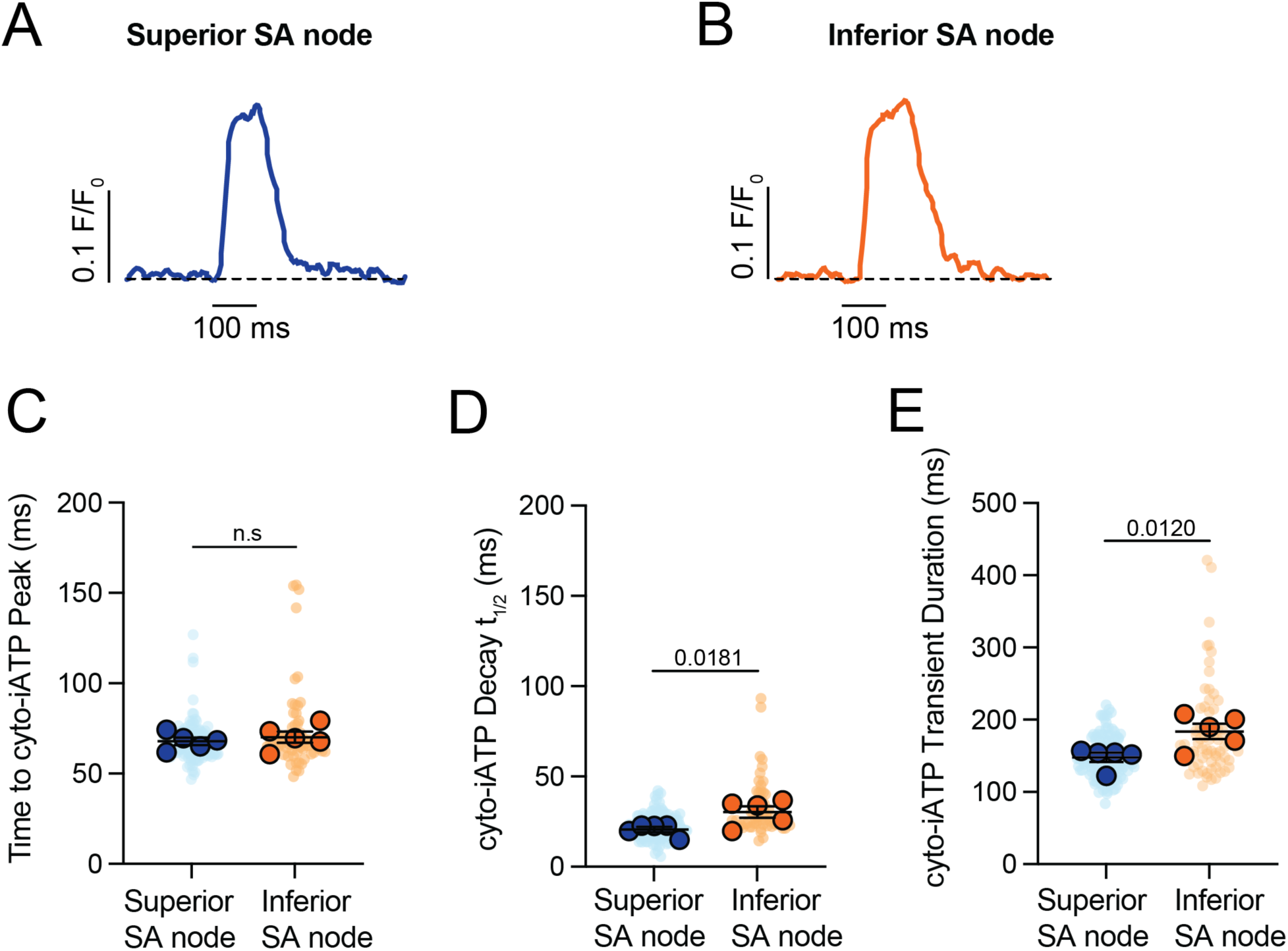
Accelerated ATP decay and shorter transient duration characterize the high-flux phenotype of the superior SA node. (**A**, **B**) Representative time-course traces of cyto-iATP transients recorded from superior (**A**) and inferior (**B**) SA node regions. Traces are normalized to peak amplitude to facilitate comparison of transient kinetics. (**C**–**E**) Summary quantification of cyto-iATP kinetic parameters (N = 5 mice per group), including time to peak (**C**), decay t_1/2_ (**D**), and total transient duration (**E**). *P*-values are indicated in the text (time to peak, *P* = 0.2777; decay t_1/2_, *P* = 0.0181; duration, *P* = 0.0120). Large circles denote per-animal means; small circles indicate individual biological replicates.

**Supplemental Figure 2.**
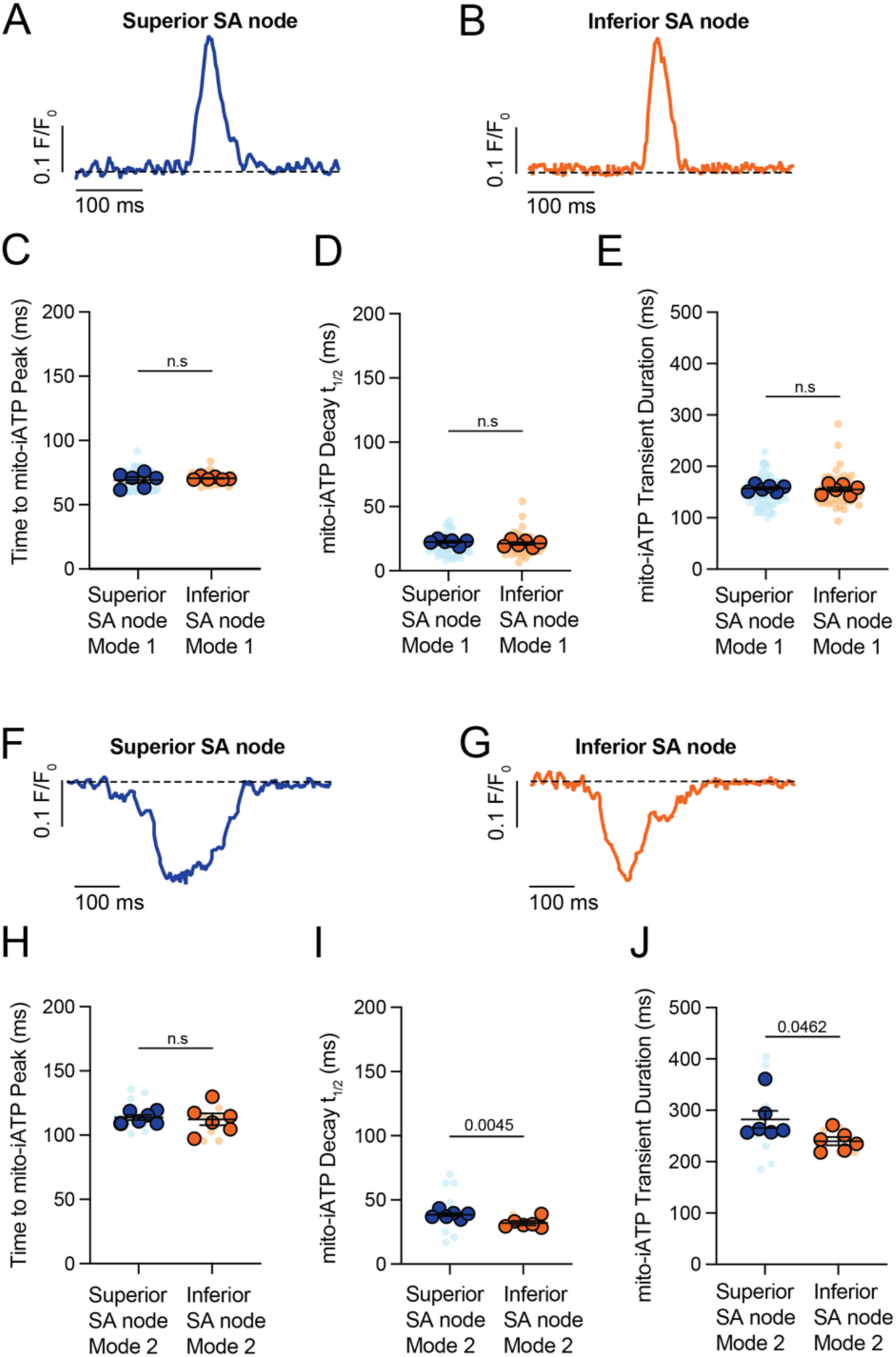
Mitochondrial ATP synthesis kinetics are regionally conserved, whereas consumption duration scales with metabolic load. (**A**–**E**) Kinetic analysis of Mode 1 (ATP production) mito-iATP transients. (**A**, **B**) Representative time-course traces from superior (**A**) and inferior (**B**) SA node regions. Traces are normalized to peak amplitude to facilitate comparison of temporal kinetics. (**C**–**E**) Summary quantification of time to peak (**C**), decay t_1/2_ (**D**), and transient duration (**E**) (N = 6 mice per group). (**F**–**J**) Kinetic analysis of Mode 2 (ATP consumption) mito-iATP transients. (**F**, **G**) Representative time-course traces from superior (**F**) and inferior (**G**) SA node regions showing negative mito-iATP deflections associated with ATP consumption. (**H**–**J**) Summary quantification of time to peak (**H**), decay t_1/2_ (**I**), and transient duration (**J**). *P*-values are shown above comparisons. Large circles denote per-animal means; small circles indicate individual biological replicates.

**Supplemental Figure 3.**
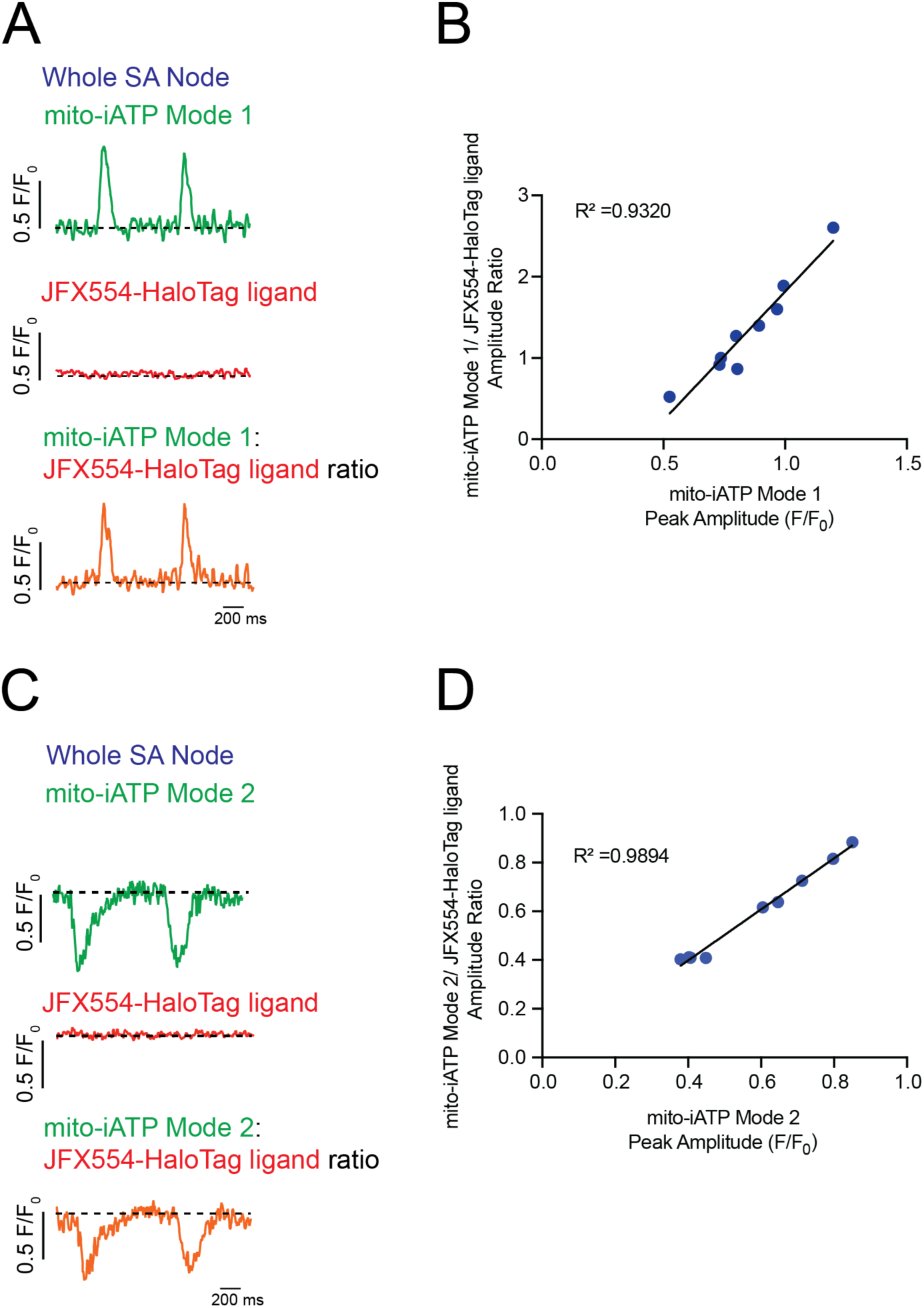
Ratiometric validation confirms that beat-to-beat mito-iATP oscillations are bona fide metabolic signals independent of motion artifacts. (**A**, **C**) Representative recordings from whole SA node preparations exhibiting Mode 1 (**A**) and Mode 2 (**C**) mito-iATP dynamics. The ATP sensor signal (green) is shown together with a spectrally distinct reference fluorophore (JFX554–HaloTag ligand, red), with the corresponding ratiometric trace overlaid (orange). (**B**, **D**) Event-level relationship between raw mito-iATP peak amplitude (F/F₀) and ratiometric amplitude for Mode 1 (**B**) and Mode 2 (**D**) events. Solid lines indicate linear fits (R^2^>0.9, both modes). Traces are displayed as normalized fluorescence (F/F₀).

**Supplemental Figure 4.**
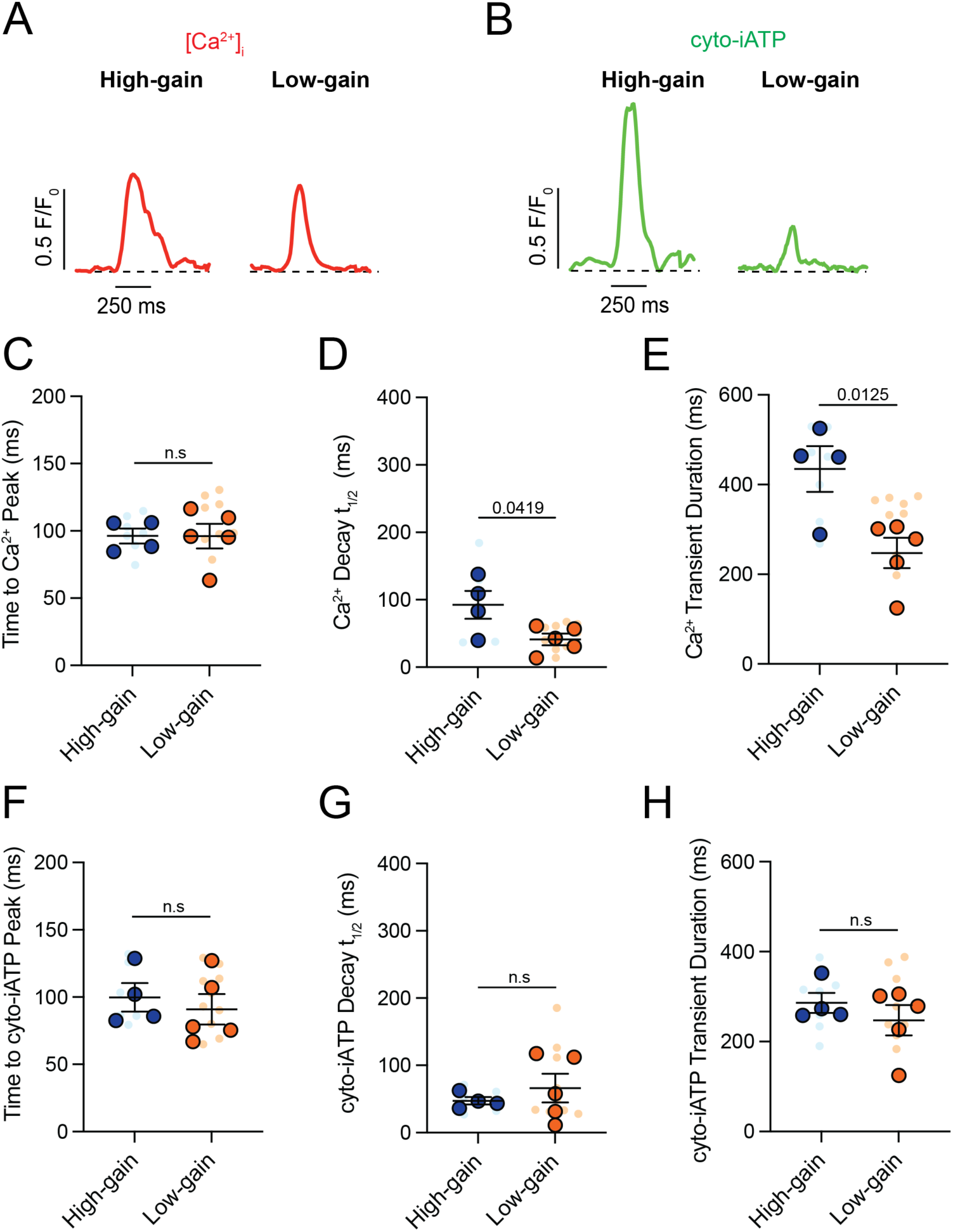
High-gain SA node myocytes exhibit prolonged intracellular Ca^2+^ transients compared to low-gain myocytes. (**A**, **B**) Representative normalized confocal line-scan recordings of intracellular Ca^2+^ (red, **A**) and cyto-iATP (green, **B**) from myocytes classified as high-gain (left) or low-gain (right). (**C**–**E**) Summary quantification of Ca^2+^ transient kinetics (high-gain, N = 4; low-gain, N = 5), including time to peak (**C**), decay t_1/2_ (**D**), and transient duration (**E**). (**F**–**H**) Summary quantification of cyto-iATP transient kinetics, including time to peak (**F**), decay constant (**G**), and transient duration (**H**). *P*-values are shown above comparisons. Large circles denote per-animal means; small circles indicate individual biological replicates.

**Supplemental Figure 5.**
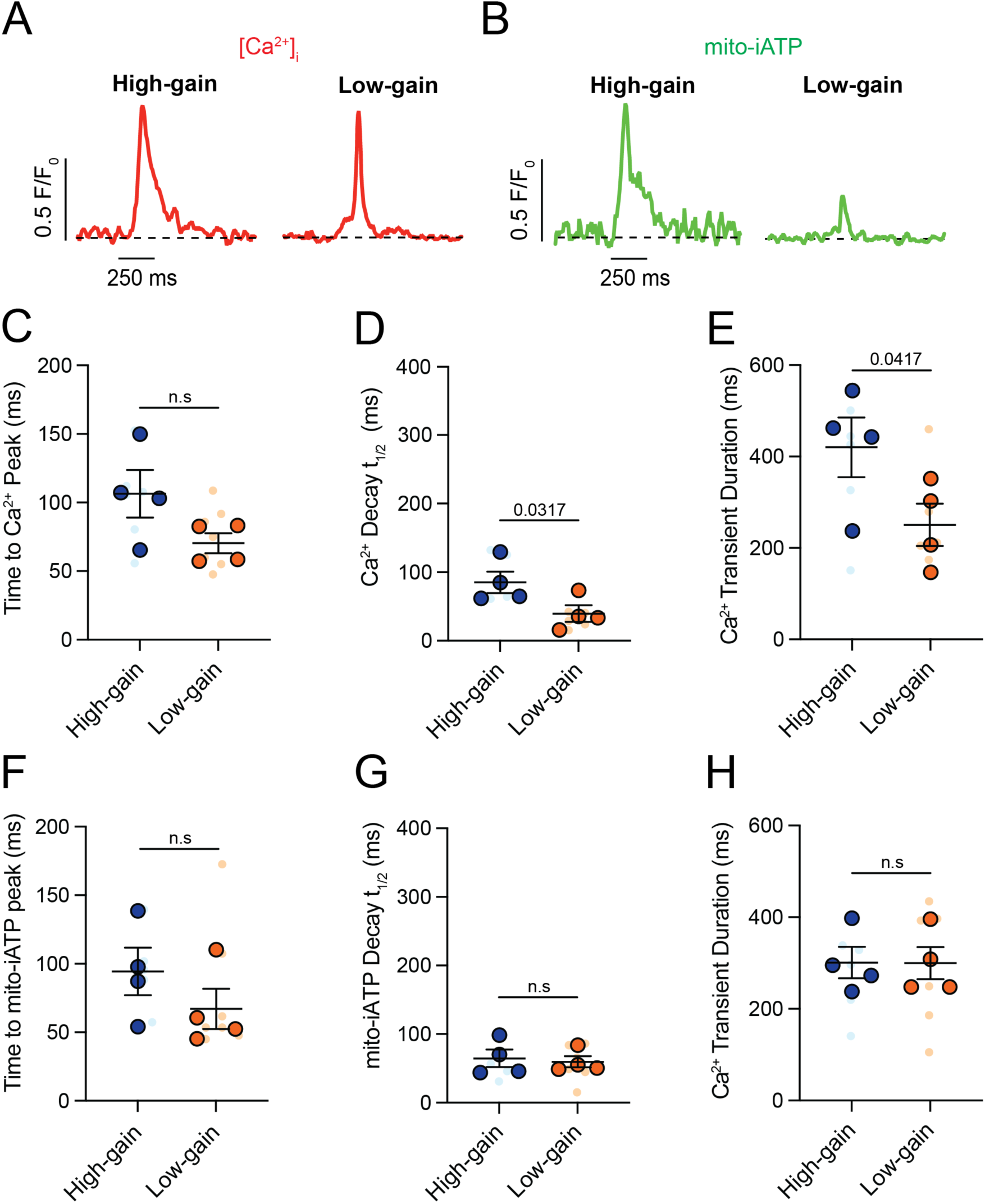
High-gain mitochondrial ATP synthesis is associated with prolonged intracellular Ca^2+^ transients. (**A**, **B**) Representative normalized confocal line-scan recordings of intracellular Ca^2+^ (red, **A**) and mito-iATP (green, **B**) from myocytes classified as high-gain (left) or low-gain (right). (**C**–**E**) Summary quantification of Ca^2+^ transient kinetics (N = 4 mice per group), including time to peak (**C**), decay t_1/2_ (**D**), and transient duration (**E**). (**F**–**H**) Summary quantification of Mode 1 mito-iATP transient kinetics, including time to peak (**F**), decay t_1/2_ (**G**), and transient duration (**H**). *P*-values are shown above comparisons. Large circles denote per-animal means; small circles indicate individual biological replicates.

**Supplemental Figure 6.**
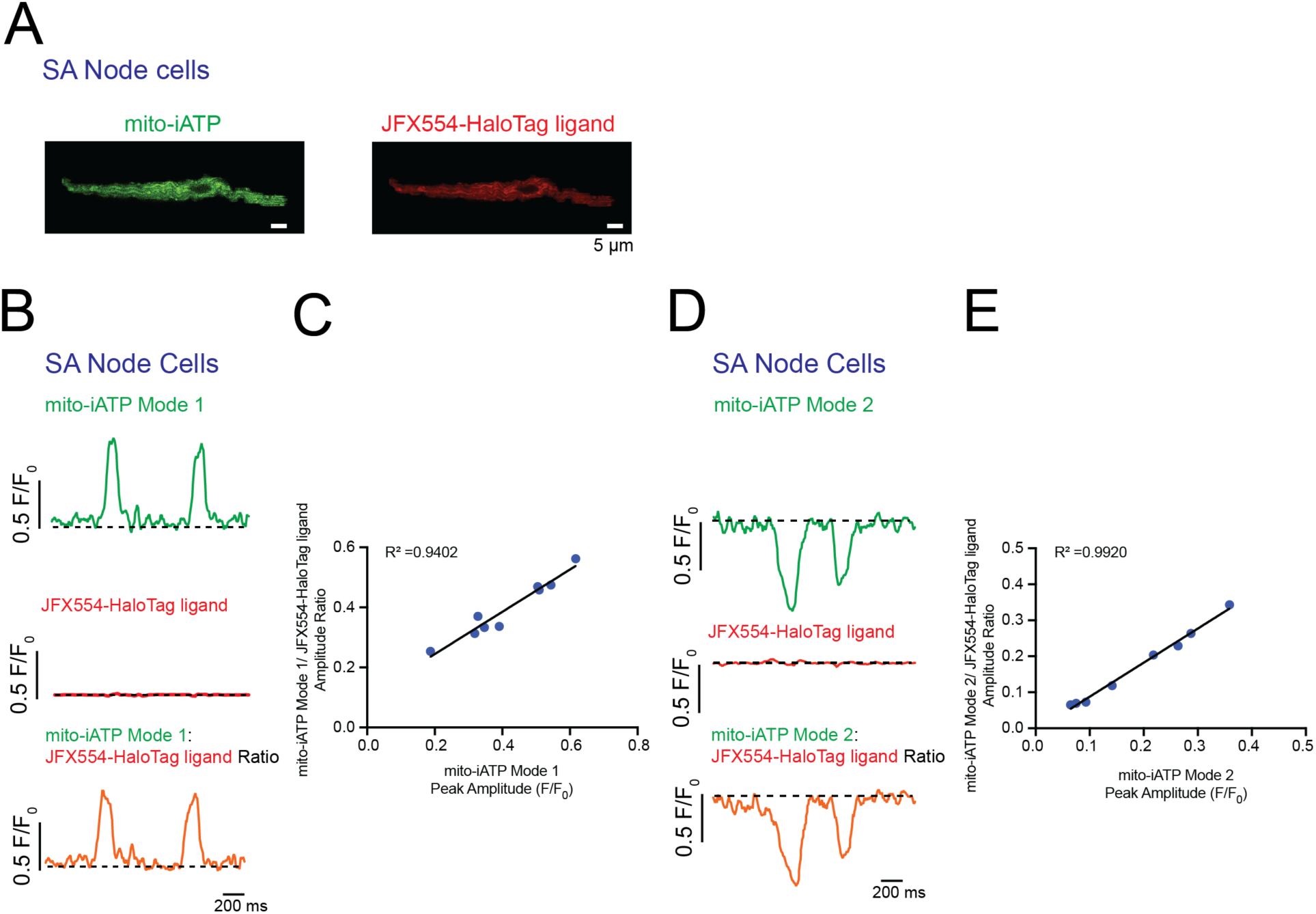
Ratiometric validation in isolated myocytes confirms that Mode 1 and Mode 2 transients reflect bona fide mitochondrial ATP dynamics. (**A**) Representative fluorescence image of an isolated SA node myocyte expressing mito-iATP (green) and labeled with the reference fluorophore JFX554–HaloTag ligand (red). Scale bar, 50 µm. (**B**, **D**) Representative single-cell recordings showing Mode 1 mito-iATP production transients (**B**) and Mode 2 mito-iATP consumption transients (**D**). The mito-iATP signal (green) is shown together with the reference fluorophore signal (red). (**C**, **E**) Event-level relationship between raw mito-iATP peak amplitude (F/F₀) and ratiometric amplitude for Mode 1 (**C**) and Mode 2 (**E**) events. Solid lines indicate linear fits. Dashed lines denote baseline fluorescence. Traces are displayed as normalized fluorescence (F/F₀). Scale bars: 0.5 F/F₀, 200 ms.

**Supplemental Figure 7.**
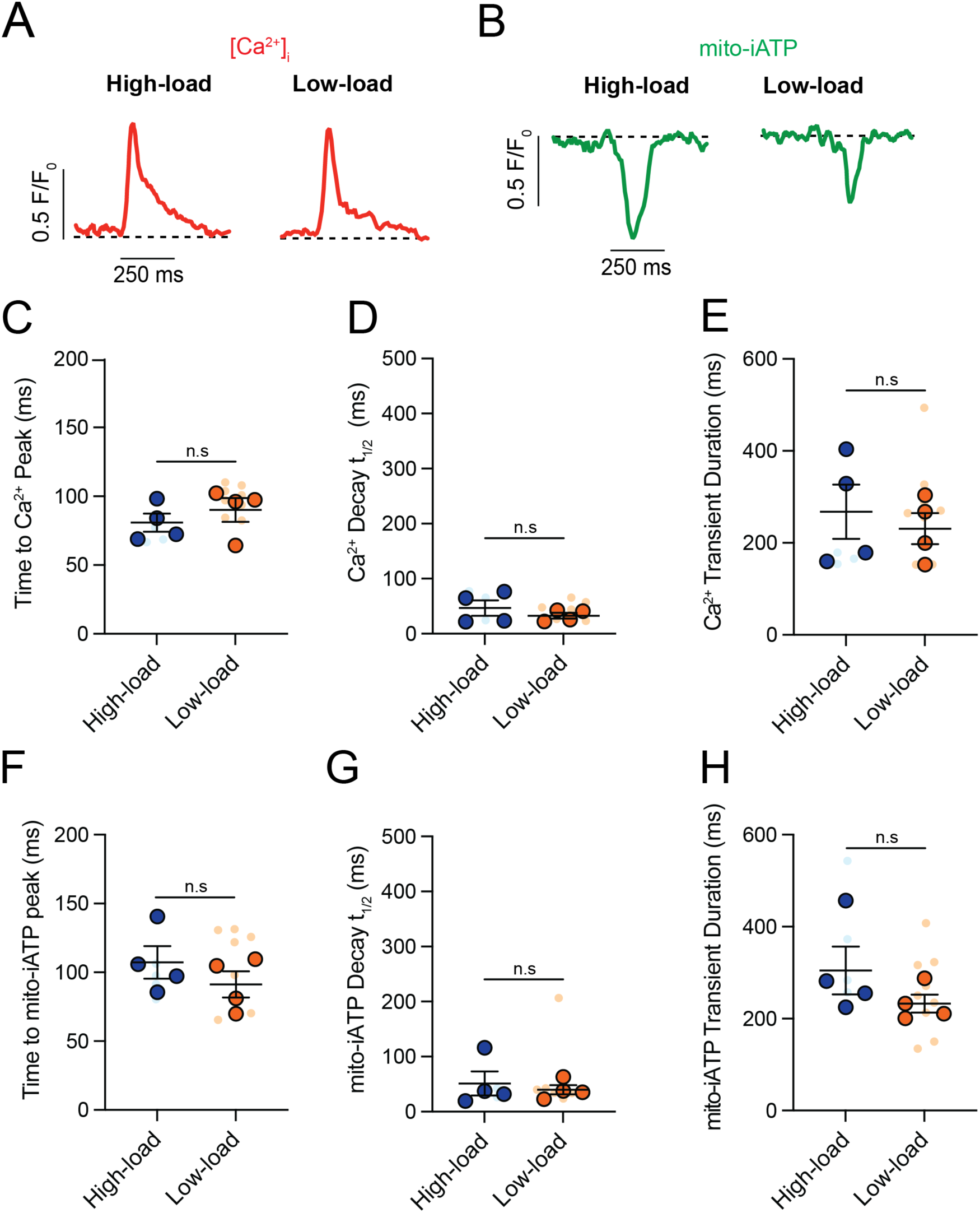
High-load mitochondrial ATP consumption is associated with intracellular Ca^2+^ release kinetics in SA node myocytes. (**A**, **B**) Representative normalized confocal line-scan recordings of intracellular Ca^2+^ (red, **A**) and Mode 2 mito-iATP signals (green, **B**) from myocytes classified as high-load (left) or low-load (right). (**C**–**E**) Summary quantification of Ca^2+^ transient kinetics (N = 4 mice per group), including time to peak (**C**), decay t_1/2_ (**D**), and transient duration (**E**). (**F**–**H**) Summary quantification of Mode 2 mito-iATP transient kinetics. *P*-values are shown above comparisons. Large circles denote per-animal means; small circles indicate individual biological replicates.

## References

1. Anumonwo, J.M., M. Delmar, A. Vinet, D.C. Michaels, and J. Jalife. 1991. Phase resetting and entrainment of pacemaker activity in single sinus nodal cells. Circ Res. 68:1138–1153.

2. Berthiaume, J.M., J.G. Kurdys, D.M. Muntean, and M.G. Rosca. 2019. Mitochondrial NAD(+)/NADH Redox State and Diabetic Cardiomyopathy. Antioxid Redox Signal. 30:375–398.

3. Bogdanov, K.Y., T.M. Vinogradova, and E.G. Lakatta. 2001. Sinoatrial nodal cell ryanodine receptor and Na(+)-Ca(2+) exchanger: molecular partners in pacemaker regulation. Circ Res. 88:1254–1258.

4. Boyett, M.R., H. Honjo, and I. Kodama. 2000. The sinoatrial node, a heterogeneous pacemaker structure. Cardiovasc Res. 47:658–687.

5. Brennan, J.A., Q. Chen, A. Gams, J. Dyavanapalli, D. Mendelowitz, W. Peng, and I.R. Efimov. 2020. Evidence of Superior and Inferior Sinoatrial Nodes in the Mammalian Heart. JACC Clin Electrophysiol. 6:1827–1840.

6. Brown, H., and D. Difrancesco. 1980. Voltage-clamp investigations of membrane currents underlying pace-maker activity in rabbit sino-atrial node. J Physiol. 308:331–351.

7. Bychkov, R., M. Juhaszova, K. Tsutsui, C. Coletta, M.D. Stern, V.A. Maltsev, and E.G. Lakatta. 2020. Synchronized Cardiac Impulses Emerge From Heterogeneous Local Calcium Signals Within and Among Cells of Pacemaker Tissue. JACC Clin Electrophysiol. 6:907–931.

8. Chen, Y., G. Csordas, C. Jowdy, T.G. Schneider, N. Csordas, W. Wang, Y. Liu, M. Kohlhaas, M. Meiser, S. Bergem, J.M. Nerbonne, G.W. Dorn, 2nd, and C. Maack. 2012. Mitofusin 2-containing mitochondrial-reticular microdomains direct rapid cardiomyocyte bioenergetic responses via interorganelle Ca(2+) crosstalk. Circ Res. 111:863–875.

9. Clancy, C.E., and L.F. Santana. 2020. Evolving Discovery of the Origin of the Heartbeat: A New Perspective on Sinus Rhythm. JACC Clin Electrophysiol. 6:932–934.

10. Csordas, G., T. Golenar, E.L. Seifert, K.J. Kamer, Y. Sancak, F. Perocchi, C. Moffat, D. Weaver, S.F. Perez, R. Bogorad, V. Koteliansky, J. Adijanto, V.K. Mootha, and G. Hajnoczky. 2013. MICU1 controls both the threshold and cooperative activation of the mitochondrial Ca(2)(+) uniporter. Cell Metab. 17:976–987.

11. Denton, R.M., and J.G. McCormack. 1990. Ca2+ as a second messenger within mitochondria of the heart and other tissues. Annu Rev Physiol. 52:451–466.

12. Denton, R.M., J.G. McCormack, and N.J. Edgell. 1980. Role of calcium ions in the regulation of intramitochondrial metabolism. Effects of Na+, Mg2+ and ruthenium red on the Ca2+-stimulated oxidation of oxoglutarate and on pyruvate dehydrogenase activity in intact rat heart mitochondria. Biochem J. 190:107–117.

13. DiFrancesco, D. 1986. Characterization of single pacemaker channels in cardiac sino-atrial node cells. Nature. 324:470–473.

14. Dorn, G.W., 2nd, M. Song, and K. Walsh. 2015. Functional implications of mitofusin 2-mediated mitochondrial-SR tethering. J Mol Cell Cardiol. 78:123–128.

15. Fenske, S., K. Hennis, R.D. Rotzer, V.F. Brox, E. Becirovic, A. Scharr, C. Gruner, T. Ziegler, V. Mehlfeld, J. Brennan, I.R. Efimov, A.G. Pauza, M. Moser, C.T. Wotjak, C. Kupatt, R. Gonner, R. Zhang, H. Zhang, X. Zong, M. Biel, and C. Wahl-Schott. 2020. cAMP-dependent regulation of HCN4 controls the tonic entrainment process in sinoatrial node pacemaker cells. Nat Commun. 11:5555.

16. Gammaitoni, L., P. Hänggi, P. Jung, and F. Marchesoni. 1998. Stochastic resonance. Reviews of Modern Physics. 70:223–287.

17. Garbincius, J.F., and J.W. Elrod. 2022. Mitochondrial calcium exchange in physiology and disease. Physiol Rev. 102:893–992.

18. Garrud, T.A.C., B. Bell, A. Mata-Daboin, D. Peixoto-Neves, D.M. Collier, J.F. Cordero-Morales, and J.H. Jaggar. 2024. WNK kinase is a vasoactive chloride sensor in endothelial cells. Proceedings of the National Academy of Sciences. 121:e2322135121.

19. Grainger, N., L. Guarina, R.H. Cudmore, and L.F. Santana. 2021. The Organization of the Sinoatrial Node Microvasculature Varies Regionally to Match Local Myocyte Excitability. Function (Oxf). 2:zqab031.

20. Guarina, L., J.T. Le, T.N. Griffith, L.F. Santana, and R.H. Cudmore. 2024. SanPy: Software for the analysis and visualization of whole-cell current-clamp recordings. Biophys J. 123:759–769.

21. Guarina, L., A.N. Moghbel, M.S. Pourhosseinzadeh, R.H. Cudmore, D. Sato, C.E. Clancy, and L.F. Santana. 2022. Biological noise is a key determinant of the reproducibility and adaptability of cardiac pacemaking and EC coupling. J Gen Physiol. 154.

22. Hanggi, P. 2002. Stochastic resonance in biology. How noise can enhance detection of weak signals and help improve biological information processing. Chemphyschem. 3:285–290.

23. Hao, Y.A., S. Lee, R.H. Roth, S. Natale, L. Gomez, J. Taxidis, P.S. O’Neill, V. Villette, J. Bradley, Z. Wang, D. Jiang, G. Zhang, M. Sheng, D. Lu, E. Boyden, I. Delvendahl, P. Golshani, M. Wernig, D.E. Feldman, N. Ji, J. Ding, T.C. Sudhof, T.R. Clandinin, and M.Z. Lin. 2024. A fast and responsive voltage indicator with enhanced sensitivity for unitary synaptic events. Neuron. 112:3680–3696 e3688.

24. Henderson, P.J., and H.A. Lardy. 1970. Bongkrekic acid. An inhibitor of the adenine nucleotide translocase of mitochondria. J Biol Chem. 245:1319–1326.

25. Huang, S., A.A. Heikal, and W.W. Webb. 2002. Two-photon fluorescence spectroscopy and microscopy of NAD(P)H and flavoprotein. Biophys J. 82:2811–2825.

26. Huser, J., L.A. Blatter, and S.L. Lipsius. 2000. Intracellular Ca2+ release contributes to automaticity in cat atrial pacemaker cells. J Physiol. 524 Pt 2:415–422.

27. Jalife, J. 1984. Mutual entrainment and electrical coupling as mechanisms for synchronous firing of rabbit sino-atrial pace-maker cells. J Physiol. 356:221–243.

28. Kamer, K.J., Z. Grabarek, and V.K. Mootha. 2017. High-affinity cooperative Ca(2+) binding by MICU1-MICU2 serves as an on-off switch for the uniporter. EMBO Rep. 18:1397–1411.

29. Kanemaru, K., J. Cranley, D. Muraro, A.M.A. Miranda, S.Y. Ho, A. Wilbrey-Clark, J. Patrick Pett, K. Polanski, L. Richardson, M. Litvinukova, N. Kumasaka, Y. Qin, Z. Jablonska, C.I. Semprich, L. Mach, M. Dabrowska, N. Richoz, L. Bolt, L. Mamanova, R. Kapuge, S.N. Barnett, S. Perera, C. Talavera-López, I. Mulas, K.T. Mahbubani, L. Tuck, L. Wang, M.M. Huang, M. Prete, S. Pritchard, J. Dark, K. Saeb-Parsy, M. Patel, M.R. Clatworthy, N. Hübner, R.A. Chowdhury, M. Noseda, and S.A. Teichmann. 2023. Spatially resolved multiomics of human cardiac niches. Nature. 619:801–810.

30. Kim, H.Y., K.Y. Lee, Y. Lu, J. Wang, L. Cui, S.J. Kim, J.M. Chung, and K. Chung. 2011. Mitochondrial Ca(2+) uptake is essential for synaptic plasticity in pain. J Neurosci. 31:12982–12991.

31. Kim, M.S., A.V. Maltsev, O. Monfredi, L.A. Maltseva, A. Wirth, M.C. Florio, K. Tsutsui, D.R. Riordon, S.P. Parsons, S. Tagirova, B.D. Ziman, M.D. Stern, E.G. Lakatta, and V.A. Maltsev. 2018. Heterogeneity of calcium clock functions in dormant, dysrhythmically and rhythmically firing single pacemaker cells isolated from SA node. Cell Calcium. 74:168–179.

32. Kirichok, Y., G. Krapivinsky, and D.E. Clapham. 2004. The mitochondrial calcium uniporter is a highly selective ion channel. Nature. 427:360–364.

33. Kühlbrandt, W. 2019. Structure and Mechanisms of F-Type ATP Synthases. Annu Rev Biochem. 88:515–549.

34. Liang, D., J. Xue, L. Geng, L. Zhou, B. Lv, Q. Zeng, K. Xiong, H. Zhou, D. Xie, F. Zhang, J. Liu, Y. Liu, L. Li, J. Yang, Z. Xue, and Y.H. Chen. 2021. Cellular and molecular landscape of mammalian sinoatrial node revealed by single-cell RNA sequencing. Nat Commun. 12:287.

35. Lobas, M.A., R. Tao, J. Nagai, M.T. Kronschlager, P.M. Borden, J.S. Marvin, L.L. Looger, and B.S. Khakh. 2019. A genetically encoded single-wavelength sensor for imaging cytosolic and cell surface ATP. Nat Commun. 10:711.

36. Lopaschuk, G.D., Q.G. Karwi, R. Tian, A.R. Wende, and E.D. Abel. 2021. Cardiac Energy Metabolism in Heart Failure. Circ Res. 128:1487–1513.

37. Mallilankaraman, K., P. Doonan, C. Cardenas, H.C. Chandramoorthy, M. Muller, R. Miller, N.E. Hoffman, R.K. Gandhirajan, J. Molgo, M.J. Birnbaum, B.S. Rothberg, D.O. Mak, J.K. Foskett, and M. Madesh. 2012. MICU1 is an essential gatekeeper for MCU-mediated mitochondrial Ca(2+) uptake that regulates cell survival. Cell. 151:630–644.

38. Mangoni, M.E., A. Traboulsie, A.L. Leoni, B. Couette, L. Marger, K. Le Quang, E. Kupfer, A. Cohen-Solal, J. Vilar, H.S. Shin, D. Escande, F. Charpentier, J. Nargeot, and P. Lory. 2006. Bradycardia and slowing of the atrioventricular conduction in mice lacking CaV3.1/alpha1G T-type calcium channels. Circ Res. 98:1422–1430.

39. Manning, D., E.J. Rivera, P. Rhana, C. Matsumoto, Z. Fong, P.N. Thai, M.F. Munoz, J.E. Contreras, S. Kim, N. Grainger, N. Chiamvimonvat, G.M. Bautista, and L.F. Santana. 2025. Microvascular Rarefaction in the Sinoatrial Node: A Potential Mechanism for Pacemaker Dysfunction in Early HFpEF. JACC Clin Electrophysiol.

40. Manning, D., E.J. Rivera, and L.F. Santana. 2024. The life cycle of a capillary: Mechanisms of angiogenesis and rarefaction in microvascular physiology and pathologies. Vascul Pharmacol. 156:107393.

41. Maravall, M., Z.F. Mainen, B.L. Sabatini, and K. Svoboda. 2000. Estimating intracellular calcium concentrations and buffering without wavelength ratioing. Biophys J. 78:2655–2667.

42. Marvin, J.S., A.C. Kokotos, M. Kumar, C. Pulido, A.N. Tkachuk, J.S. Yao, T.A. Brown, and T.A. Ryan. 2024. iATPSnFR2: A high-dynamic-range fluorescent sensor for monitoring intracellular ATP. Proc Natl Acad Sci U S A. 121:e2314604121.

43. Matlib, M.A., Z. Zhou, S. Knight, S. Ahmed, K.M. Choi, J. Krause-Bauer, R. Phillips, R. Altschuld, Y. Katsube, N. Sperelakis, and D.M. Bers. 1998. Oxygen-bridged dinuclear ruthenium amine complex specifically inhibits Ca2+ uptake into mitochondria in vitro and in situ in single cardiac myocytes. J Biol Chem. 273:10223–10231.

44. McCormack, J.G., A.P. Halestrap, and R.M. Denton. 1990. Role of calcium ions in regulation of mammalian intramitochondrial metabolism. Physiol Rev. 70:391–425.

45. Monfredi, O., K. Tsutsui, B. Ziman, M.D. Stern, E.G. Lakatta, and V.A. Maltsev. 2018. Electrophysiological heterogeneity of pacemaker cells in the rabbit intercaval region, including the SA node: insights from recording multiple ion currents in each cell. Am J Physiol Heart Circ Physiol. 314:H403–H414.

46. Okamura, A., I.K. He, M. Wang, A.V. Malsev, A.V. Maltsev, M.D. Stern, E.G. Lakatta, and V.A. Maltsev. 2024. Cardiac Pacemaker Cells Harness Stochastic Resonance to Ensure Fail-Safe Operation at Low Rates Bordering on Sinus Arrest. bioRxiv.

47. Rhana, P., C. Matsumoto, Z. Fong, A.D. Costa, S.G. Del Villar, R.E. Dixon, and L.F. Santana. 2024. Fueling the heartbeat: Dynamic regulation of intracellular ATP during excitation-contraction coupling in ventricular myocytes. Proc Natl Acad Sci U S A. 121:e2318535121.

48. Rhana, P., C. Matsumoto, and L.F. Santana. 2025. Demonstration of Beat-to-Beat, On-Demand ATP Synthesis in Ventricular Myocytes Reveals Sex-Specific Mitochondrial and Cytosolic Dynamics. bioRxiv.

49. Rowe, G.C., A. Jiang, and Z. Arany. 2010. PGC-1 coactivators in cardiac development and disease. Circ Res. 107:825–838.

50. Ruprecht, J.J., M.S. King, T. Zogg, A.A. Aleksandrova, E. Pardon, P.G. Crichton, J. Steyaert, and E.R.S. Kunji. 2019. The Molecular Mechanism of Transport by the Mitochondrial ADP/ATP Carrier. Cell. 176:435–447 e415.

51. Saddik, M., and G.D. Lopaschuk. 1991. Myocardial triglyceride turnover and contribution to energy substrate utilization in isolated working rat hearts. J Biol Chem. 266:8162–8170.

52. Santana, L.F., and S. Earley. In press. Energetic Microdomains and the Vascular Control of Neuronal and Muscle Excitability: Toward a Unified Model The Jounal of Physiology.

53. Scarpulla, R.C. 2011. Metabolic control of mitochondrial biogenesis through the PGC-1 family regulatory network. Biochim Biophys Acta. 1813:1269–1278.

54. Tagirova Sirenko, S., K. Tsutsui, K.V. Tarasov, D. Yang, A.N. Wirth, V.A. Maltsev, B.D. Ziman, Y. Yaniv, and E.G. Lakatta. 2021. Self-Similar Synchronization of Calcium and Membrane Potential Transitions During Action Potential Cycles Predict Heart Rate Across Species. JACC Clin Electrophysiol. 7:1331–1344.

55. Walker, J.E. 2013. The ATP synthase: the understood, the uncertain and the unknown. Biochem Soc Trans. 41:1–16.

56. Wiesenfeld, K., and F. Moss. 1995. Stochastic resonance and the benefits of noise: from ice ages to crayfish and SQUIDs. Nature. 373:33–36.

57. Wisneski, J.A., W.C. Stanley, R.A. Neese, and E.W. Gertz. 1990. Effects of acute hyperglycemia on myocardial glycolytic activity in humans. J Clin Invest. 85:1648–1656.

58. Yamamoto, M., H. Honjo, R. Niwa, and I. Kodama. 1998. Low-frequency extracellular potentials recorded from the sinoatrial node. Cardiovasc Res. 39:360–372.

59. Yaniv, Y., H.A. Spurgeon, B.D. Ziman, A.E. Lyashkov, and E.G. Lakatta. 2013. Mechanisms that match ATP supply to demand in cardiac pacemaker cells during high ATP demand. Am J Physiol Heart Circ Physiol. 304:H1428–1438.

60. Zhou, B., and R. Tian. 2018. Mitochondrial dysfunction in pathophysiology of heart failure. J Clin Invest. 128:3716–3726.

